# Tau in the brain interstitial fluid is fragmented and seeding-competent

**DOI:** 10.1101/2020.07.15.205724

**Authors:** Erica Barini, Gudrun Plotzky, Yulia Mordashova, Jonas Hoppe, Esther Rodriguez-Correa, Sonja Julier, Florie LePrieult, Ina Mairhofer, Mario Mezler, Sandra Biesinger, Miroslav Cik, Marcus W Meinhardt, Ebru Ercan-Herbst, Dagmar E. Ehrnhoefer, Andreas Striebinger, Karen Bodie, Corinna Klein, Laura Gasparini, Kerstin Schlegel

**Author notes:** Equal contributions (Laura Gasparini, Kerstin Schlegel). Department of Molecular Neuroimaging and Institute of Psychopharmacology, Central Institute of Mental Health, Medical Faculty Mannheim, University of Heidelberg, 68159 Germany (Marcus W Meinhardt).

## Abstract

In Alzheimer disease, Tau pathology is thought to propagate from cell to cell throughout interconnected brain areas. However, the forms of Tau released into the brain interstitial fluid (ISF) in vivo during the development of Tauopathy and their pathological relevance remain unclear. Combining in vivo microdialysis and biochemical analysis, we find that in Tau transgenic mice, human Tau (hTau) present in brain ISF is truncated and comprises at least 10 distinct fragments spanning the entire Tau protein. The fragmentation pattern is similar across different Tau transgenic models, pathological stages and brain areas. ISF hTau concentration decreases during Tauopathy progression, while its phosphorylation increases. ISF from mice with established Tauopathy induces Tau aggregation in HEK293-Tau biosensor cells. Notably, immunodepletion of ISF phosphorylated Tau, but not Tau fragments, significantly reduces its ability to seed Tau aggregation and only a fraction of Tau, separated by ultracentrifugation, is seeding competent. These results indicate that ISF seeding competence is driven by a small subset of Tau, which potentially contribute to the propagation of Tau pathology.

**Highlights:**

- In interstitial fluid (ISF) of transgenic mice, Tau comprises >10 distinct fragments
- ISF Tau decreases with Tauopathy progression, while its phosphorylation increases
- Only ISF from mice with established Tauopathy is seeding competent in vitro
- Removal of phospho-Tau reduces ISF seeding competence
- ISF seeding competence is driven by less soluble, aggregated and phosphorylated Tau

**Graphical abstract:** 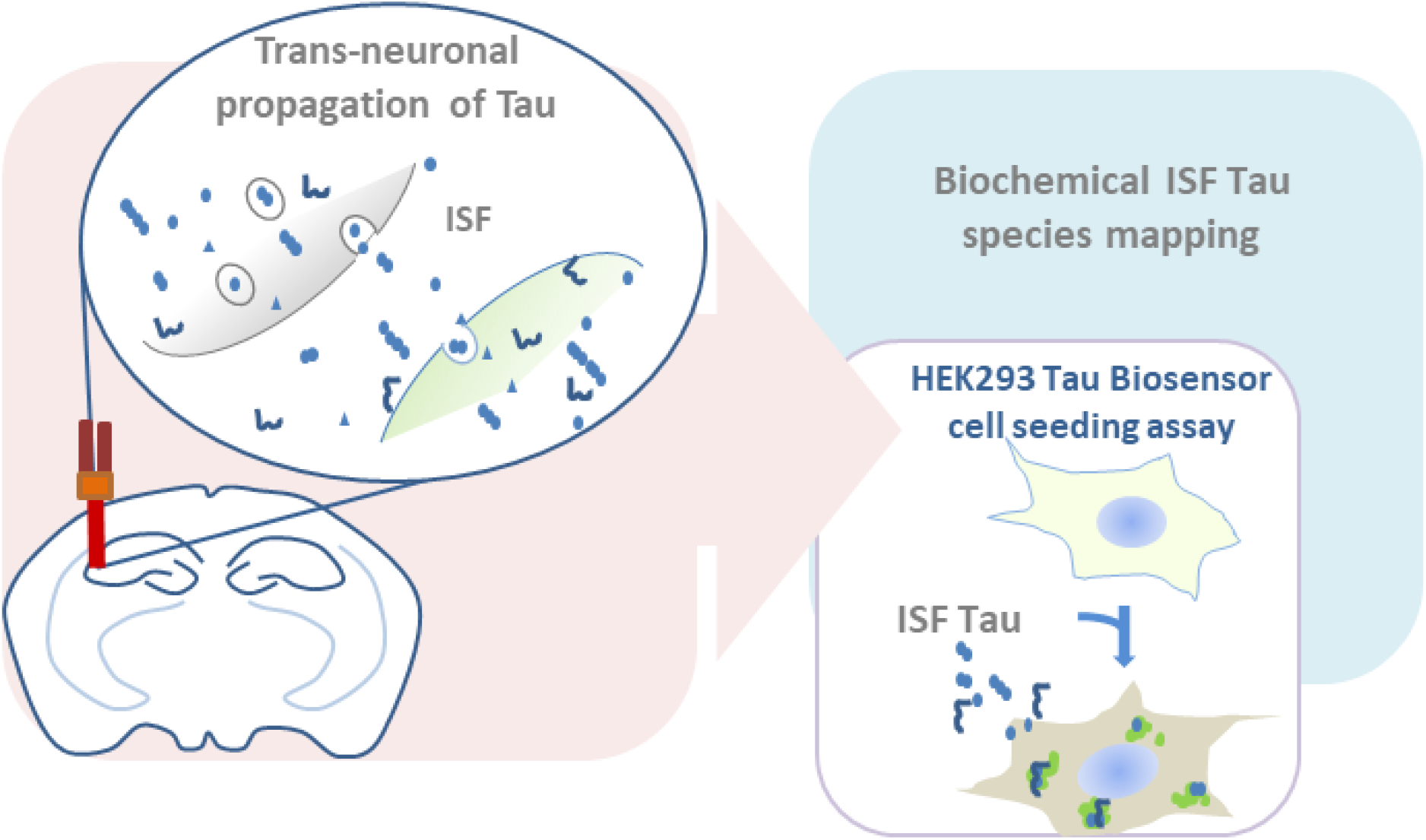

## 1. Introduction

Intracellular inclusions of the microtubule-associated protein Tau are neuropathological hallmarks of a subset of neurodegenerative diseases called Tauopathies, including Alzheimer disease (AD), corticobasal degeneration (CBD), progressive parasupranuclear palsy (PSP) and frontotemporal dementia (FTD) (Lee et al., 2001). In AD, Tau pathology develops in a hierarchical manner, progressively spreading throughout anatomically-interconnected brain regions, from the entorhinal cortex and hippocampus at early Braak stages I-II, to the occipito-temporal and insular cortex at Braak stages III-IV, to the entire neocortex at the advanced stages Braak V-VI (Braak and Braak, 1991).

In recent years, in vitro and in vivo experimental models consistently indicate that pathological Tau species propagate across interconnected neurons. In cultured primary neurons, Tau is released into the extracellular space in an activity-dependent manner (Chai et al., 2012, Pooler et al., 2013) through unconventional secretion (Chai et al., 2012, Pooler et al., 2013), and is subsequently taken up by synaptically-connected neurons, where it nucleates and seeds further aggregation (Wu et al., 2016). Using push-pull microdialysis to sample brain interstitial fluid (ISF) in living, freely-moving PS19 Tau transgenic mice, it has been demonstrated that full-length Tau can be detected in the extracellular compartment and ISF Tau levels decrease during pathology progression (Yamada et al., 2011) and increase upon K^+^-evoked neuronal depolarization (Yamada et al., 2014).

The nature of pathological, seeding-competent, propagating Tau species remains unknown. Ex vivo fractionation studies of extracts from human brain indicate that only soluble aggregates such as trimers, oligomers or short fibrils likely comprising hyperphosphorylated and nitrated Tau are internalized by neurons and induce Tau aggregation in vitro and in vivo (Jackson et al., 2016, Mirbaha et al., 2015). In these studies, the discovery of such soluble, seeding-competent species relied upon analysis of tissue homogenates, which include both intracellular and extracellular Tau species. It therefore remains unclear whether the identified Tau species are indeed those released in the ISF and subsequently taken up by connected neurons. Studies in cultured neurons in vitro have reported that full-length and truncated Tau is secreted in free form (Sato et al., 2018, Bright et al., 2015) or included in small vesicles called exosomes (Asai et al., 2015), indicating that extracellular Tau may comprise a variety of species. It has been also shown that ISF Tau sampled from rTg4510 mice is taken up by primary neurons in culture (Takeda et al., 2015) and by HEK293-Tau biosensor cells, where it induces Tau aggregate formation (Takeda et al., 2016), but the ISF Tau species triggering this effect has not been determined.

So far, only little data is available on ISF Tau composition (Yamada et al., 2011, Bright et al., 2015). Studies by independent groups in two Tau transgenic mouse lines reported apparently inconsistent findings. Yamada and colleagues (2011) found that in ISF of PS19 Tau transgenic mice, full-length monomeric Tau is the most abundant Tau species. Instead, in ISF from JNPL3 P301L Tau mice, Bright et al. (2015) mainly found a truncated Tau form of approximately 25 kDa as detected by an N-terminal Tau antibody. Thus, the identity of ISF Tau species and their pathological relevance for Tauopathy progression remains unclear.

Here, to define the composition of ISF Tau and its seeding competence, we applied state-of-the-art in vivo microdialysis to sample ISF from freely moving Tau transgenic mice during Tau pathology development. We found that ISF Tau is largely truncated and includes at least 10 distinct fragments spanning the entire protein. The fragmentation pattern is similar across three different Tau transgenic mouse strains (i.e. PS19, rTg4510, Thy1.P301Stau), pathological stages and brain areas. The overall concentration of human Tau (hTau) is inversely related to its phosphorylation state and ISF seeding competence. In fact, hTau levels decrease during Tauopathy progression, while phosphorylation on selected Tau residues increases. Moreover, only ISF from mice with established pathology induces Tau aggregation in HEK293-Tau biosensor cells. We also found that immunodepletion of soluble Tau fragments does not reduce ISF seeding competence, whereas removal of phosphorylated Tau species significantly decreases the ability to induce Tau aggregation in vitro. Notably, ISF seeding-competent Tau species could be sedimented by ultracentrifugation: the supernatant containing all soluble Tau fragments does not have any seeding propensity, while less soluble Tau species in the pellet are seeding competent. These results indicate that only a fraction of ISF Tau species account for ISF seeding competence and might have a role in the pathological propagation of Tau throughout the brain.

## 2. Materials and Methods

### 2.1. Reagents and antibodies

Antibodies used for western blots and immunoprecipitation (IP) are listed in Table 1. The following reagents were used: 50 mM Tris-HCl pH7.5 (Alpha Diagnostics Inc.), phosphate-buffered saline (PBS; Lonza, BioWhittaker #BE17-517Q), bovine serum albumin (BSA, Serva #11926.03 or Blocker™ 10% BSA for western blot washing buffer, ThermoFisher, Catalogue No. 37525), 1,4-dithiothreitol (DTT, Sigma #10197777001), Laemmli Sample Buffer (LDS, 4x Laemmli Sample Buffer BIORAD #1610747), Dimethyl pimelimidate dihydrochloride (DMP, Thermo Fisher cat 21667), Tween-20 (Sigma #P1379-100ML). We used the primary mouse monoclonal and rabbit polyclonal antibodies listed in Table 1. Tau12 antibody was biotinylated with Pierce™ Antibody Biotinylation Kit for IP (90407). Secondary biotinylated anti-mouse (Jackson #115-065-166) and anti-rabbit (Jackson #111-065-144) antibodies were used for detection.

**Table 1:** Antibodies used for western blot and IP.

| Antibody Name | Species | Epitope | Supplier | Cat. n° | IP | ELISA | WB | Western Blot concentration |
| --- | --- | --- | --- | --- | --- | --- | --- | --- |
| Tau12 | mouse | 6-18 | BioLegend | 806501 | ✓ | ✓ |  |  |
| Tau12<br>(in-house biotinylation) | mouse | 6-18 | BioLegend | 806501 |  |  | ✓ | 1µg/mL |
| HT7 | mouse | 159-163 | Thermo Fisher | MN1000 | ✓ | ✓ |  |  |
| HT7-Biotin | mouse | 159-163 | Thermo Fisher | MN1000B |  |  | ✓ | 0.2µg/mL |
| BT2 | mouse | 194-198 | Thermo Fisher | MN1010 | ✓ |  |  |  |
| BT2-Biotin | mouse | 194-198 | Thermo Fisher | MN1010 B |  |  | ✓ | 0.1µg/mL |
| AT100 | mouse | pT212/pS214 | Thermo Fisher | MN1060 |  | ✓ |  |  |
| pT181 | rabbit | pT181 | Thermo Fisher | 701530 |  | ✓ |  |  |
| pS198 | rabbit | pS198 | Abcam | ab79540 |  | ✓ |  |  |
| pS199 | rabbit | pS199 | Thermo Fisher | 701054 |  | ✓ |  |  |
| pT231 | rabbit | pT231 | Thermo Fisher | 701056 |  | ✓ |  |  |
| pS416 | mouse | pS416 | Abcam | ab119391 |  | ✓ |  |  |
| Tau 5 | mouse | 210-241 | Calbiochem | 577801 |  |  | ✓ | 1µg/mL |
| AB64193 | rabbit | p262/262 | Abcam | AB64193 | ✓ |  | ✓ | 0.4µg/mL |
| HJ8.7 | mouse/human | 117-122 | Holtzman lab | (Yanamandra et al., 2017) |  |  | ✓ | 0.2 µg/mL |
| HJ8.3 | mouse/human | 405-411 | Holtzman lab |  |  |  | ✓ | 1µg/mL |
| HJ9.1 | mouse/human | 385-401 | Holtzman lab |  | ✓ |  |  |  |

### 2.2. Experimental animal models

Heterozygous PS19 mice (licensed from University of Pennsylvania, Philadelphia, PA, USA; (Yoshiyama et al., 2007), homozygous Thy1.P301Stau (licensed from the Medical Research Council, UK) (Allen et al., 2002), heterozygous rTg4510 (licensed from the Mayo Clinic, Jacksonville Florida, USA; (Santacruz, 2005 #595)(Santacruz et al., 2005), Tau knockout (TauKO; JAX stock number: 004779) (Tucker et al., 2001) and wild -type mice were used. All transgenic mice were bred for Abbvie by Charles River Laboratories (Sulzfeld, Germany). Animal health and comfort were veterinary-controlled. Mice were in a temperature- and humidity-controlled room with a 12:12 hour dark/light cycle with *ad libitum* access to water and food. All animal experiments were performed in full compliance with the Principles of Laboratory Animal Care (NIH publication No. 86-23, revised 1985) in an AAALAC-accredited program where veterinary care and oversight was provided to ensure appropriate animal care. All animal studies were approved by the government of Rhineland Palatinate (Landesuntersuchungsamt Koblenz, Germany) and conducted in accordance with the directive 2010/63/EU of the European Parliament and of the Council on the protection of animals used for scientific purposes, the ordinance on the protection of animals used for experimental or scientific purposes (German implementation of EU directive 2010/63; BGBl. I S. 3125, 3126), the Commission Recommendation 2007/526/EC on guidelines for the accommodation and care of animals used for experimental and other scientific purposes and the German Animal Welfare Act (BGBl. I S. 1206, 1313) amended by Art. 1 G from 17 December 2018 I 2586.

PS19 mice express 1N4R human P301S mutant Tau under the control of PrP promoter (Yoshiyama et al., 2007). rTg4510 mice express 0N4R Tau under the CaMKIIα promoter. rTg4510 mice progressively accumulate sarkosyl-insoluble hTau and AT100 immunoreactive (AT100+) inclusions throughout the forebrain from 4 months of age (Santacruz et al., 2005). Thy1.P301Stau mice express 0N4R human P301S mutant Tau under the control of the mouse neuronal Thy.1 promoter. They progressively develop widespread filamentous inclusions made of hyperphosphorylated Tau throughout the cerebral cortex, brain stem, spinal cord and retina, accompanied by neuronal and motor dysfunction and paraparesis (Scattoni et al., 2010, Allen et al., 2002, Gasparini et al., 2011). Tau knockouts are homozygous for the EGFP coding sequence inserted into the first exon of the *Mapt* gene, disrupting its expression (Tucker et al., 2001).

### 2.3. Aseptic surgery of mice

All surgeries were performed under aseptic conditions using sevoflurane anesthesia. Guide cannulas PEG-4 (EICOM) or cOFM guides (Joanneum GD-2-2, Joanneum Research Forschungsgesellschaft mbH) targeted the left and right hippocampus (AP-3, ML-/+3.1, DV-2.7 for EICOM and AP-3, ML-/+3.1, DV-4 for cOFM) of PS19, rTg4510 or TauKO mice. In Thy1.P301Stau mice, guides were embedded in the right hippocampus (AP-3, ML-3.1, DV-2.7) and in the left cortex (AP 2.2, ML 1.5, DV-0.93). In selected experiments, in rTg4510 mice, cOFM guide cannulas (Joanneum GD-2-2) and PEG-4 (EICOM) were implanted in the right (AP-3.2, ML-3.1, DV-4) and the left hippocampus (AP-3, ML +3.1, DV-2.7), respectively. The skull was cemented and mice were kept at 37°C to recover post-surgery until awake.

### 2.4. *In vivo* Tau microdialysis

Two weeks after surgery, mice were briefly anesthetized using isoflurane and 1MDa cut-off EICOM or cOFM probes connected to the peristaltic pump operating in push-pull mode were inserted into the respective guide cannulas. Mice were placed in a cage coupled with a BASi Stand-Alone Raturn System. ISF sampling ran up to 48 h after probe insertion in awake, freely-moving mice. Usually, 8 to 12 mice underwent microdialysis simultaneously and ISF was collected via a refrigerated fraction collector (Microbiotech MAB80) with a sampling time of 60 min. ISF vials (polystyrene, Microbiotech) were either left empty or filled with 6 μl of a mix of post-translational modification (PTM) inhibitors containing the following reagents: phosphatase inhibitors (PhosSTOP, 04906845001, Roche), protease inhibitors (cOmplete Protease Inhibitor Cocktail, Roche, 11697498001), 500 mM IOX1 (Sigma-Aldrich, SML0067), 2 mM Daminozide (Sigma-Aldrich, 45418), 10 mM Trichostatin A (Sigma-Aldrich, T8552), 5 mM Nicotinamide (Sigma-Aldrich, 72340), 10 mM Pargyline HCl (Sigma-Aldrich, P8013), 1 mM Thiamet G (Sigma-Aldrich, SML0244). The flow was constantly monitored throughout the experiment either by weighing samples or online flow sensors (Sensirion, SLG-0150). After microdialysis, brains were analyzed by an experienced pathologist by histology to ensure proper probe placement and evaluate tissue inflammatory reactions.

For the investigation of Tau levels under basal conditions or upon K+ stimulation, the setup was prepared as follows: FEB-Tubing and Santopren tubing were cut and connected to the push and pull system. Tubes were flushed with 5% Pluronic Acid (Sigma, P2443) overnight and with 1% Cold Sterilant (VWR #148-0062) for 30 min. The perfusate solution contained artificial cerebrospinal fluid (aCSF), which was prepared using the following reagents: 0.52 mM CaCl_2_ x 2 H2O (Sigma, C3306-100G), 48.8 mM NaCl (Merck, 1064041000), 0.16 mM NaH_2_PO_4_ x H2O (Merck, 1063460500), 0.48 mM MgSO_4_ x 7 H2O (Merck, 1058860500), 1.2 mM KCl (Merck, 1049361000), 10.4 mM NaHCO_3_ (Merck, 1063291000). The aCSF solution was mixed with 0.2% BSA (Sigma, A8577-10ML), filtered through a 0.1μm membrane and flushed at 1 μl/min for at least 30 min before animal connection. For K^+^-induced neuronal depolarization, perfusate solution containing 100 mM KCl was unilaterally perfused, while plain perfusate solution (vehicle) was used in the contralateral hemisphere. Before K+ stimulation, up to 4 samples were collected and analyzed to define hTau levels at baseline.

For comparison between ISF sampled with 2-mm 1 MDa cut-off EICOM probes and cOFM (Joanneum COFM-2-2) probes, the experiment was performed as follows: FEB tubing was used for the left hemisphere, where the PEG-4 guide (EICOM) was implanted, and PVC tubing (OFM-PP2-LT-500/300, assembled by Joanneum) was used for the right hemisphere, where the cOFM guides were implanted. Before animal connection, tubes were flushed with 5% Pluronic Acid, with 1% Cold Sterilant and with aCSF and 0.2% BSA, as previously described.

For all other microdialysis experiments, cOFM probes were used and the setup was prepared as follows: 1% Cold Sterilant (VWR #148-0062) was flushed for 30 min and washed out with Ampuwa sterile water (Sigma, P2443-250G). aCSF containing 0.2% BSA was then flushed at 1 μl/min for at least 30 min before animal connection.

### 2.5. Immunoprecipitation and immunodepletion

IP was performed using Dynabeads Protein G (Invitrogen, #1003D). 30 mg of magnetic beads were mixed with 50 μg of antibody overnight at 4°C and covalently coupled with 25 mM DMP in 0.2 M Triethanolamine-HCl pH 8.2 for 30 min. The reaction was neutralized with 50 mM Tris-HCL pH 7.5. 50 μl of ISF with phosphatases and proteases inhibitors was added to 10 μl of bead-antibody complex and incubated at 4°C overnight shaking at 1250 rpm. After washing with PBS plus 0.05% Tween-20 (PBST), samples were boiled in 1X LDS with 50 mM DTT and for 5 min at 98°C. Samples were then stored at −20°C until use.

For immunodepletion studies, Protein G Dynabeads were freshly prepared under sterile conditions. All solutions were sterile filtered with 0.2 μm filters (Midisart^®^ 2000). Coupling of antibodies to beads was performed as described above. Beads were washed in PBS without detergents. 20 μl of Dynabead-antibody complex was incubated with 100 μl of ISF at 4°C shaking overnight at 1250 rpm. Dynabead-antibody-Tau complexes were isolated using a magnetic holder and the supernatant was used for further assays. 50 μl of ISF input was always kept at 4°C until use as a reference.

### 2.6. Western blot

Equal volumes of ISF were boiled at 98°C for 5 min in 50 mM of DTT and 1X LDS and separated by electrophoresis on 4-12% Bis-Tris Precast Gels (BioRad #3450124). Proteins were transferred onto 0.2 μm PVDF membranes (Biorad #1704157) using Trans Blot Turbo (BioRad). Membranes were boiled in PBS pH 7.4 for 20 min, blocked overnight at 4°C with 5% BSA in PBS-T, incubated with antibodies for 2 h at room temperature (RT), and rinsed in 1X Tris-buffered saline pH 7.4 (TBS, Biorad #1706435) containing 0.05% Tween-20 (TBST) followed by 0.01% BSA (Blocker™ 10% BSA) in PBS. Membranes were then incubated for 1 h at room temperature with secondary biotinylated antibodies or with the ABC kit (Cat. no. 32020, Thermo Fisher). SuperSignal West Femto (Thermo Scientific Cat n. 34095) or Clarity™ (BioRad # 1705061) was used for signal detection. Signal was captured using Chemidoc (Biorad) and densitometric analysis was performed using ImageLab (BioRad).

### 2.7. hTau, phospho-Tau and aggregated Tau analyses

In all ISF samples, levels of hTau were routinely measured using the MSD V-Plex Human Total Tau Kit (Meso Scale Diagnostics, LLC, K151LAG) according to the manufacturer’s instructions. Briefly, plates were blocked using 150 μl of Diluent35 per well and incubated at RT for 1 h at 800 rpm. 50 μl of sample/well was used. The plate was incubated for 1 h at RT before adding 25 μl/well of detection antibody solution and washing. Ready Buffer T was then added, and plates were analyzed with an MSD imager 6000.

Levels of Tau phosphorylated at T181, S198, S199, T231 and S461 in the ISF were measured with MSD V-Plex Human Total Tau plates to capture Tau, where the detection was done by using sulfo-tagged phospho-sepecific antibodies. Briefly, ISF was diluted 1:4 with Diluent35 and added to the plates, followed by 1 h incubation at room temperature. For detection, antibodies detecting the phospho-sites were sulfo-tagged following the manufacturer’s manual (Cat. No. R91AO-1). Plates were analyzed on the MSD Quickplex platform. Aggregated Tau was measured by ELISA using Tau-12 for both capture and detection (Ercan-Herbst et al., 2019). Levels of pT212/pS214 Tau were measured by ELISA using AT100 for capture and HT7 for detection. A standard curve was plotted based on recombinant phospho-Tau produced in Sf9 insect cells. Plates were analyzed using a Versa Max Plate Reader (Molecular Devices) at OD 450 nm.

### 2.8. HEK293-Tau biosensor cells seeding assay

Seeding assay experiments were performed using HEK293-Tau biosensor cells stably expressing the hTau-RD-P301S-YFP construct (Holmes and Diamond, 2014) as previously described (Takeda et al., 2016) with minor modifications. In brief, HEK293-Tau biosensor cells were plated at defined cell density into 96-well clear bottom microtiter plates and cultured for 24 h at 37°C and 8% CO_2_. The day of the experiment, 2000 cells/well were plated in a 384-well plate (Corning #354663). Serial dilutions (1:3, 1:6 and 1:9 or 1:10 and 1:100) of ISF samples were prepared using OPTIMEM+ Glutamax-1 (Gibco #51985-026) and kept on ice until lipofection. 40 μM recombinant Tau seeds in PBS were serially diluted to obtain a standard curve ranging from 0.2 to 50 pM. 2% Lipofectamine (Lipofectamin 2000 Reagent, Invitrogen #11668-019) in OPTIMEM+ Glutamax-1 (Gibco #51985-026) was kept 120 min at RT before mixing 1:1 with samples and adding to cells. 48 h post-lipofection, cells were fixed for 30 min at RT in 4% of PFA containing 4% sucrose. Cell nuclei were stained by Hoechst 33342 and imaged with the Cellomics Array Scan HCA. The number of cells with one or more YFP aggregates was counted using an automated algorithm and dose-response curves were fitted using a sigmoidal dose-response model with variable slope using Graph Pad PRISM 8.0 software. Seeding capability of samples was calculated based on the reference Tau standard curve.

### 2.9. Immunohistochemistry

Immunohistochemical analysis on PS19 mouse brain was performed on frozen, free-floating sections. Immunohistochemical analysis of rTg4510 and Thy1.P301Stau mouse brain was performed using paraffin-embedded sections. In brief, animals were anesthetized with Ketamin/Xylazin and transcardially perfused with PBS. Hemibrains were immersion-fixed for 24 h at RT. For paraffin sections, the tissue was transferred to 70% ethanol. Tissue was trimmed and samples dehydrated using a standard EtOH series followed by xylene and paraffin embedding (ASP300, Leica). 4 μm paraffin sections were prepared and processed on a Leica BOND Rx automated stainer using citrate-buffer based heat-induced antigen retrieval solution (Epitope Retrieval Solution 1 Bond, Leica, Germany) and DAB-based detection (Leica, Germany). For frozen, free-floating sections, tissue was cryoprotected after immersion fixation and 40 μm sections were prepared on a cryo-microtome (Reichert-Jung). Free-floating sections were quenched with 0.3% H_2_O_2_-methanol PBS solution and permeabilized in Tris-buffered saline mixed with 0.05% Tween-20 in in Tris-buffered saline and DAB-based detection (VectaStain, Biozol Germany). Tau pathology was detected using AT8 (MN1020; ThermoFisher Scientific) or AT100 anti-pS212/pT214 Tau conformational antibody (MN1060; ThermoFisher Scientific) (Augustinack et al., 2002). Images were collected with a 3DHistec P1000 scanner or Zeiss AxioScan scanner.

### 2.10. Ultracentrifugation of ISF

To fractionate ISF Tau species based on their density, pools of different ISF fractions prepared under sterile conditions were transferred into ultracentrifugation tubes (Beckman, #357448) and spun at 150,000 x g for 16 h at 4°C in a TLA 55 rotor on a Beckmann ultracentrifuge. The supernatant was collected and used for the seeding assay. ISF pellets were resuspended in PBS. Pellets, inputs and supernatants were vortexed for 1 min in PBS or TBS, depending on downstream analysis, were sonicated (2 rounds at 40% amplitude for 5 s each) and then used for the seeding assay or biochemical analyses.

### 2.11. Statistical methods

Experimental data were analyzed using parametric and non-parametric methods depending on the underlying distribution of the data. Student’s t-test, paired t-test, Welch’s test, one-way or two-way analysis of variances (ANOVA) were followed by post-hoc tests presuming normal or log-normal underlying distributions or data were further transformed (see Supplementary Table 1 for details). To explore the underlying distribution of the data, Shapiro-Wilk tests were performed to assess normality. If needed, data were transformed onto a logarithmic scale or otherwise and the homogeneity of variances was assessed using Leven’s test. One-way or two-way ANOVAs were followed by test contrast, Tukey’s HSD or Dunnett’s post-hoc test depending on the objective or design of the experiment. Non-parametric tests (Wilcoxon test, Fisher-Pitman test; Dunn’s test) were used when the assumption of normality was violated. For IP experiments, Taylor expansion was used to obtain an approximation of the mean of the ratio of the immunodepleted supernatant vs. input (both measured in quadruplicates). A summary of all statistical assumptions and tests is presented in Supplementary Table 1. Statistical analyses were carried out with Graph Pad Prism (Version 7.00), JMP 13.1.0 (SAS institute) Software and R studio Version 1.1.453. Exact p-values are reported in the figure legends. In the figure plots, for simplicity, statistical significance is indicated by * and ** for p<0.05 and p<0.01, respectively.

## 3. Results

### 3.1. ISF levels of hTau decrease with pathology progression in Tau transgenic mice

In initial experiments, we investigated how ISF levels of total hTau vary during the development of Tau pathology in three different Tau transgenic mouse models (PS19, rTg4510 and Thy1.P301Stau). These models differ in their transgene expression pattern (ubiquitous vs neuronal), features of Tau aggregates (hyperphosphorylated and/or argyrophilic) and temporal development of Tau inclusions.

As they age, PS19 mice accumulate phosphorylated Tau throughout the cortex but develop only scarce, short, argyrophilic Tau filaments (Yoshiyama et al., 2007). At 10 months of age, there are only a few sparse neurons immunoreactive for the phosphorylation- and conformation-dependent AT100 antibody (AT100+) and positively stained by thioflavin S (Fig. S1A-B), present throughout the brain. Using push-pull microdialysis through conventional 1MDa cut-off probes, we sampled ISF in awake, freely-moving PS19 mice at 4 and 9 months of age and measured hTau levels using the human Tau MSD assay. Consistent with previous data (Yamada et al., 2011), ISF hTau levels were 3.7-fold higher at 4 months of age (84.6 ± 17.8 ng/mL) compared to 9 months of age (22.7 ± 5.5 ng/mL; Fig. 1A).

**Figure 1.**
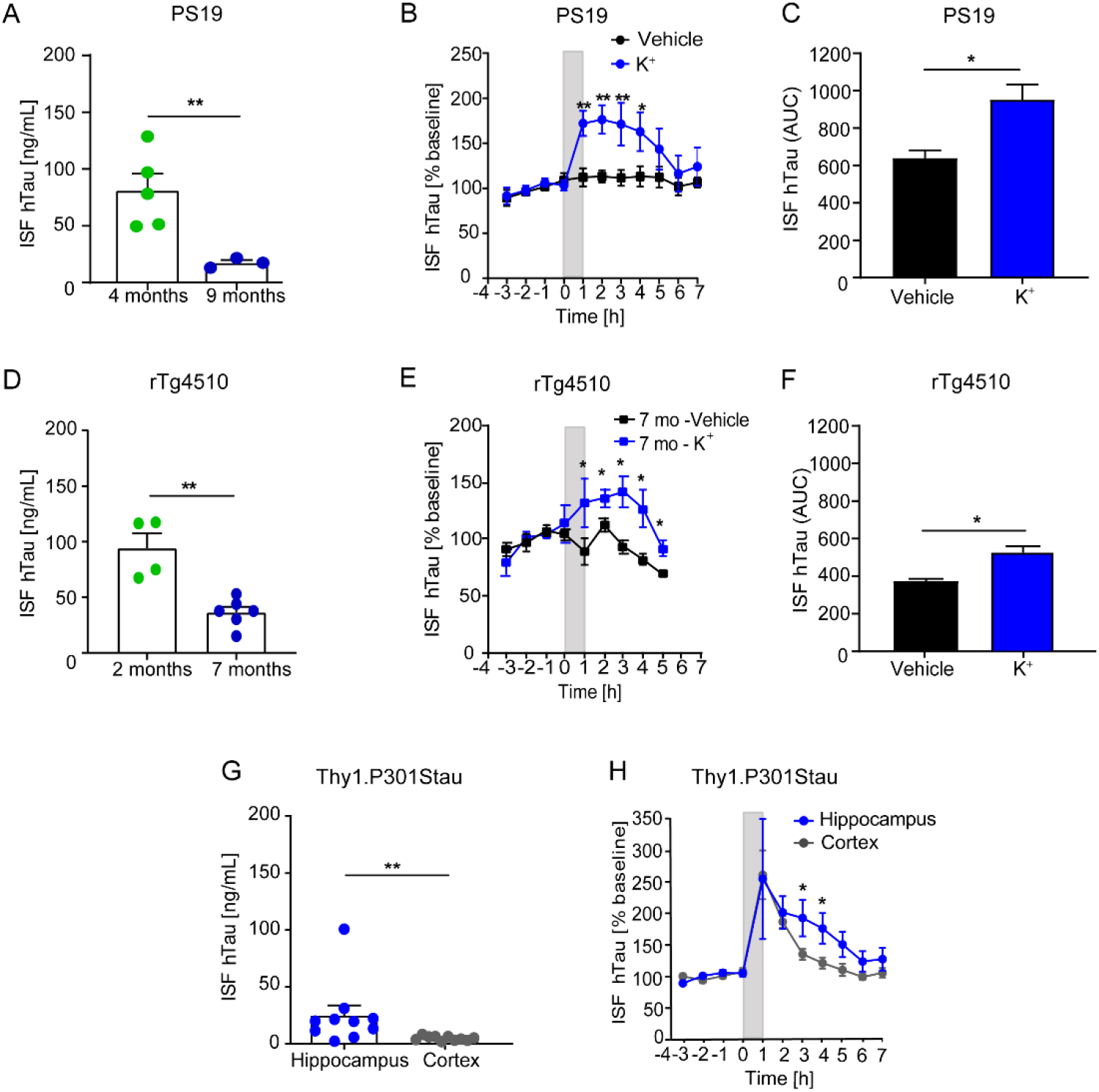
Extracellular hTau decreases during progression of Tau pathology and is increased by K^+^-evoked neuronal depolarization. In Tau transgenic mice, ISF was sampled in basal conditions during and after perfusion with 100 mM K^+^ or vehicle. (A) Basal levels of ISF hTau in 4-month-old (n=5) and 9-month-old (n=3) PS19 mice (p=0.0014). (B) ISF hTau in K^+^- or vehicle-perfused 9-month-old mice. Statistical significance: 1h, p=0.0078; 2h, p=0.0053; 3h, p=0.0080; 4h, p=0.0236; 5h, p=0.1436; 6h, p=0.4968; 7h, p=0.3981. (C) Mean area under the curve (AUC) of ISF hTau after K^+^- or vehicle-perfusion in PS19 mice (p=0.0296). (D) Basal levels of ISF hTau in rTg4510 mice at 7 months of age (n=6; p=0.0042). (E) ISF hTau in K^+^- or vehicle-perfused 7-month-old rTg4510 mice. Statistical significance: 1h, p=0.0453; 2h, p=0.0374; 3h, p=0.0256; 4h, p=0.0195; 5h, p=0.0154. (F) Mean AUC of ISF hTau after K^+^- or vehicle-perfusion in rTg4510 mice (p=0.0178). (G) Basal levels of ISF hTau in the cortex and hippocampus of 5-month-old Thy1.P301Stau mice (n=7; p=0.0021). (H) ISF hTau in K^+^-hippocampus and cortex of 5-month-old Thy1.P301Stau mice (n=10). Statistical significance: 1h, p=0.7390; 2h, p=0.8381; 3h, p=0.0414; 4h, p=0.0322; 5h, p=0.0656; 6h, p=0.2814; 7h, p=0.3865. In A, D and G, bars represent the mean + SEM and data points indicate values from single mice. In B, E and H, ISF hTau is expressed as mean % of baseline ± SEM. In C and F, bars represent the mean AUC + SEM. \**p* < 0.05, \*\**p* < 0.01

Since the presence or absence of insoluble intracellular Tau inclusions may differentially affect extracellular hTau levels, we sampled ISF from rTg4510 mice, which progressively develop abundant sarkosyl-insoluble Tau filaments and AT100+ neuronal inclusions throughout the forebrain from 4 months of age (Fig. S1C-D) (Santacruz et al., 2005). In ISF from 7-month-old rTg4510 mice compared to that of 2-month-old, levels of hTau were significantly reduced by 2.3-fold (Fig. 1D). To investigate whether the reduction of ISF Tau is simply a reflection of aging, we examined the levels of endogenous mouse Tau (mTau) in the ISF of C57 Bl/6J wild type and rTg4510 mice. In C57 Bl/6J wild type mice, mTau ISF levels were comparable at 2 (2.4 ± 0.3 ng/mL) and 10 months of age (3.3 ± 0.6 ng/mL, p=0.3, Student t-test). Consistently, mTau ISF levels in rTg4510 were comparable at 2 and 7 months of age (2 months: 4.5 ± 0.5 ng/mL; 7 months: 2.9 ± 0.5 ng/mL; p=0.2, Student t-test).

To determine the relationship between extracellular hTau levels and the abundance of Tau inclusions in different brain areas, we next used Thy1.P301Stau transgenic mice. In this model, at 5 months of age, AT100+ inclusions are abundant in the cortex, but sparse in the hippocampus (Fig. S1E-F). To sample ISF from these brain regions, each mouse was implanted with one probe in the hippocampus and one in the contralateral cortex. At 5 months of age, basal levels of ISF hTau in the cortex were 5-fold lower than in the hippocampus (Fig. 1G). There was a similar difference in hTau levels in ISF from 2-month-old Thy1.P301Stau transgenic mice, where cortical ISF contained 3.2-fold less Tau than hippocampal ISF (Fig. S2B).

Overall, these data suggest that basal levels of extracellular hTau are inversely related to the degree of Tau accumulation as mature AT100+ Tau inclusions and/or phospho-Tau aggregates.

### 3.2. The release of hTau in ISF is similarly stimulated by potassium depolarization at all Tauopathy stages

It has been previously reported that ISF Tau levels are enhanced upon stimulation with high potassium (K^+^) in PS19 mice at 3-5 months of age (Yamada et al., 2014). To understand how ISF hTau dynamics vary in a later stage of Tau pathology, we measured ISF hTau levels upon K^+^ stimulation in PS19 Tau transgenic mice at 9 months of age. 100 mM K^+^ was unilaterally delivered by reverse microdialysis for 1 h, with vehicle administered through the contralateral probe. In 9-month-old PS19 mice, ISF hTau levels significantly increased over baseline at 1 h after K^+^ stimulation, plateaued for about 3 h and returned to baseline 5 h later (Fig. 1B). The calculated area under the curve (AUC) was significantly different between the vehicle-perfused and the K^+^-stimulated hemispheres (Fig. 1C).

We next extended the analysis to rTg4510 and Thy1-P301Stau transgenic mice at different stages of Tau pathology and in different brain areas. We first investigated hTau dynamics in rTg4510 mice devoid of (2-month-old) or with (7-month-old) AT100+ inclusions. While vehicle-treated contralateral levels of ISF hTau remained unchanged, unilateral K^+^-stimulation significantly increased ISF hTau over baseline at 2 (Fig. S2A) and 7 months of age (Fig. 1G). The AUC was significantly higher after K^+^-stimulation as compared to vehicle in 7-month-old mice (Fig. 1F). Cortex and hippocampus of Thy1.P301Stau transgenic mice showed similar effects immediately after K^+^ stimulation: in both brain areas, high K^+^ increased hTau levels to comparable extents at 2 and 5 months of age (Fig. 1H; Fig. S2C). At 5 months, hTau levels in the hippocampus seem to decline slower than in the cortex (Fig. 1H).

These results indicate that K^+^-evoked neuronal depolarization enhances hTau release in ISF and the effect is independent of basal levels of extracellular hTau, Tau pathology stage and/or brain area.

### 3.3. Tau in ISF of Tau transgenic mice is present mainly as truncated forms

To investigate the composition of ISF Tau species, we immunoprecipitated Tau from pooled samples of ISF from 2- and 7-month-old rTg4510 mice using a mix of Tau antibodies targeting epitopes along the entire Tau sequence (Tau-12, HT7, AB64193 and HJ9.1; Table 1) and analyzed the pull-down by western blot using the individual antibodies. ISF from Tau knockouts (TauKO) was used in all analyses to control for band specificity (Fig. S3B and data not shown). The N-terminus antibody Tau12 mainly recognized four bands of approximately 33, 27, 20 and 17 kDa apparent molecular weight (Fig. 2A; bands A, B, C and D, respectively). Mid-domain antibodies HT7 and Tau5 detected fragments A-C and one additional fragment E of about 16 kDa (Fig. 2A, mid-panels). The Tau repeat region antibody AB64193 recognized band A and the two additional bands F and G running below 10 kDa (Fig. 2A, right panel). The pattern of Tau fragments was similar in ISF from rTg4510 (Fig 2A), Thy1.P301Stau (Fig. 2B) and PS19 mice (Fig. S3A) at all ages and across different brain regions (i.e. cortex and hippocampus; Fig. 2B). To detect the presence of fragments containing the C-terminal domain of Tau, we analyzed ISF by western blot using the antibody HJ8.3 recognizing the Tau epitope at amino acid 405-411. Three bands of about 12, 14 and 16 kDa were detected in ISF from both rTg4510 and Thy1.P301Stau mice (Fig. S3B).

**Figure 2.**
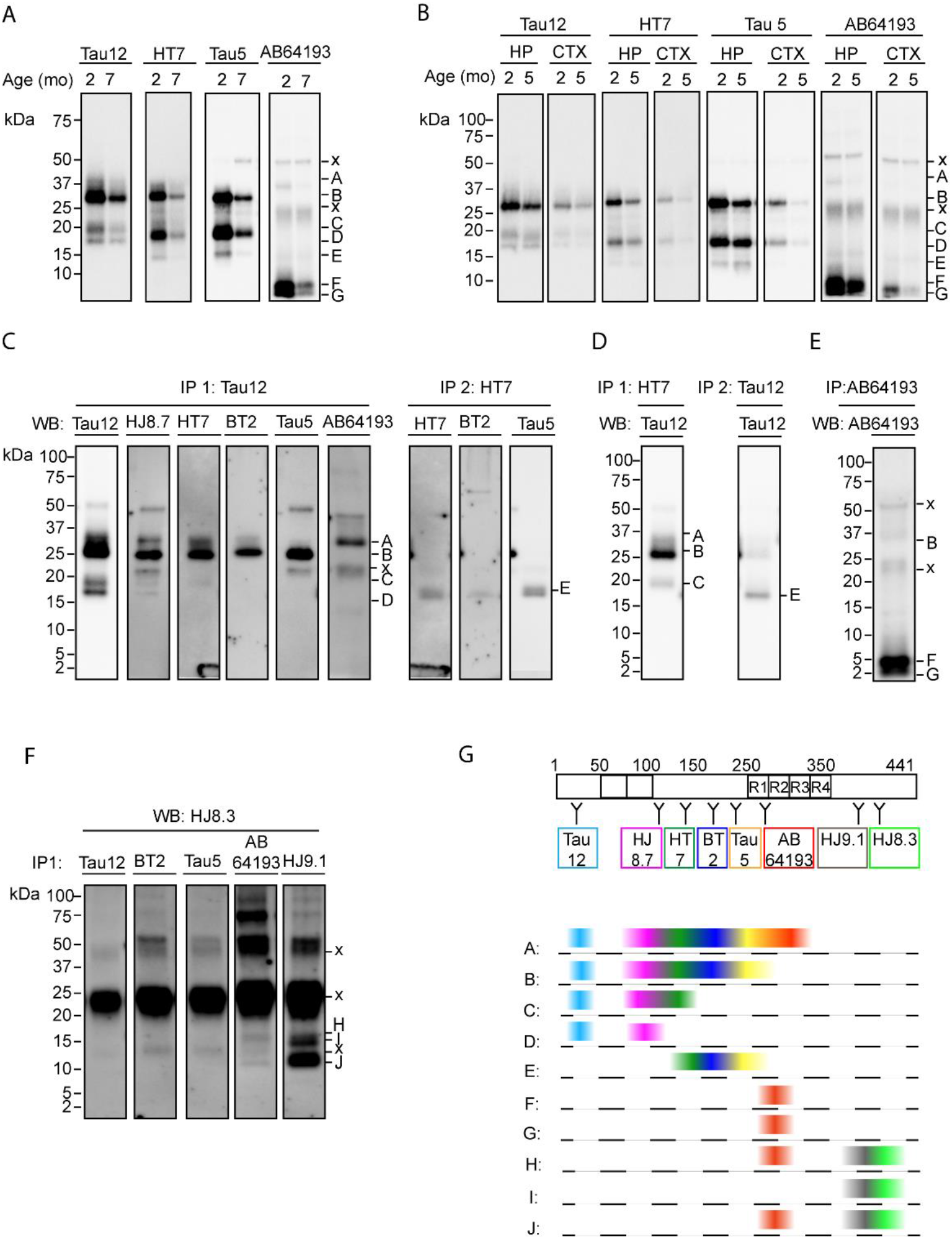
ISF Tau is fragmented into a similar pattern across transgenic mice, pathology stages and brain areas. (A-B) ISF pooled from different animals was immunoprecipitated with a mix of Tau antibodies including Tau12, HT7, AB64193 and HJ9.1 and analyzed by western blot. (A) Analysis of 2-month and 7-month-old rTg4510 ISF. (B) Analysis of ISF from 2-month-old and 5-month-old hippocampus and cortex of Thy1.P301Stau mice. (C-F) Tau fragments were analyzed by sequential immunoprecipitation (IP) and western blot in pooled ISF from rTg4510 mice. Antibodies used for IP1, IP2 and western blot are indicated in the figure. Across the panels, bands A-J represent Tau fragments; bands labeled ‘x’ are non-specific. (G) Schematic of Tau fragments mapped according to antibody epitopes and color-coded: Tau 12, light blue; HJ8.7, pink; HT7, dark green; BT2, blue; Tau5, yellow; AB64193, red; HJ9.1, grey; HJ8.3, light green.

Next, we systematically mapped the Tau fragments by sequential IP of pooled ISF from 2- and 7-month-old rTg4510 mice. ISF from age-matched TauKO mice was used to control for non-specific bands. Different Tau antibodies were used in two sequential IP rounds (IP1 and IP2) and subsequent western blot analysis (Fig. 2C; see Table 1 for antibody epitopes). Four fragments (fragments A-D) were immunoprecipitated by both N-terminus Tau12 and mid-domain HJ8.7 antibodies. Fragment A (~33 kDa) was additionally detected by antibodies HT7, BT2, and AB64193, indicating that this fragment stretches from the N-terminus until ~aa262 included. Fragment B (~27 kDa) included both the Tau12 and Tau5 epitopes (Fig. 2C, IP1). Fragment C (~20 kDa) extended from the N-terminus to the HT7 epitope and was not detected by Tau5 (Fig. 2C, IP1). Fragment D (~ 17 kDa) was the shortest of the N-terminal fragments and could only be recognized by Tau12 and HJ8.7. A weak band corresponding to full-length Tau could only be detected after IP with Tau12 and at longer exposure times (Fig. S3C). In IP2, following Tau12 IP1, Fragment E (~16 kDa), which was not recognized by Tau12, was detected by HT7, BT2 and Tau5 (Fig. 2C, IP2), indicating that this fragment contains the Tau mid-domain but not the N-terminus. Reversing the order of the antibodies (HT7 for IP1 and Tau12 for IP2) yielded similar results (Fig. 2D). Fragments A-C captured by HT7 were detected by Tau12, while fragment D was only detected after IP using Tau12 and did not contain the HT7 epitope (Fig. 2D). Two additional small bands were detected only after IP with AB64193 followed by detection with the same antibody (Fig. 2E), suggesting that such fragments span several amino acids around residue ~262. Lastly, Tau C-terminal fragments H and J were identified after IP with AB64193 and HJ9.1 and detection with HJ8.3 antibodies (Fig. 2F). Bands H and J were not detected after immunoprecipitation with Tau12, BT2 and Tau5, suggesting that these two fragments span from the first repeat R1 to part of Tau C-terminal domain. Band I was only detected by HJ8.3 after IP with HJ9.1, suggesting that this fragment contains only the C-terminal region of Tau.

In summary, these results show that the majority of Tau present in ISF is truncated and comprises at least 10 different fragments (Fig. 2G) spanning the entire Tau sequence. In Tau transgenic mice, the fragmentation pattern remains constant across pathology stages and brain areas.

### 3.4. ISF from rTg4510 induces Tau aggregation in HEK293-Tau biosensor cells

We next explored whether ISF Tau is capable of inducing Tau aggregation in HEK293-Tau biosensor cells (Kfoury et al., 2012). Previous findings showed that pooled ISF sampled from 6-13-month-old rTg4510 with 1MDa cut-off probes induced Tau aggregation in HEK293-Tau biosensor cells (Takeda et al., 2016). To understand seeding competence in more detail, we sampled ISF using either 1MDa cut-off microdialysis probes or cerebral open flow microperfusion (cOFM) probes, which enables collection of ISF without molecular size restriction. Six-month-old rTg4510 mice and control littermates were implanted with a 1MDa cut-off probe in the left hippocampus or a cOFM probe in the right hippocampus (Fig. 3A) and ISF was sampled from both hemispheres simultaneously. Seeding competency was subsequently tested using the HEK293-Tau biosensor cell assay as previously described (Holmes and Diamond, 2014). The presence of Tau aggregates was analyzed 48 h after transfection by fluorescence microscopy. Consistent with previous findings (Takeda et al., 2016), ISF sampled from rTg4510 mice using 1MDa cut-off probes significantly induced Tau aggregation in HEK293-Tau biosensor cells, while ISF from control littermates (Fig. 3B) or perfusate solution (aCSF with 0.2% BSA) did not (data not shown). In fact, Tau aggregates were increased about 4.8-fold by ISF from rTg4510 compared to control littermates (Fig. 3B). Interestingly, the seeding competency of ISF sampled by cOFM was greater than ISF sampled through 1MDa cut-off probes. Indeed, compared to ISF from control littermates, undiluted cOFM ISF from rTg4510 mice increased Tau aggregates in HEK293-Tau biosensor cells about 27.6-fold (Fig. 3B). ISF dilution resulted in a significant, progressive reduction in Tau aggregates, indicating that seeding competence depends on ISF concentration (rTg4510, cOFM, percentage of cells with Tau aggregates ± SEM: Undiluted, 30.6 ± 9.7%, 10X dilution, 12.2 ± 4.5%, 100X dilution, 3.0 ± 0.7%. rTg4510, 1MDa cut-off probe: Undiluted, 5.4 ± 1.0%, 10X dilution, 3.2 ± 0.6%, 100X dilution, 2.3 ± 0.5%).

**Figure 3.**
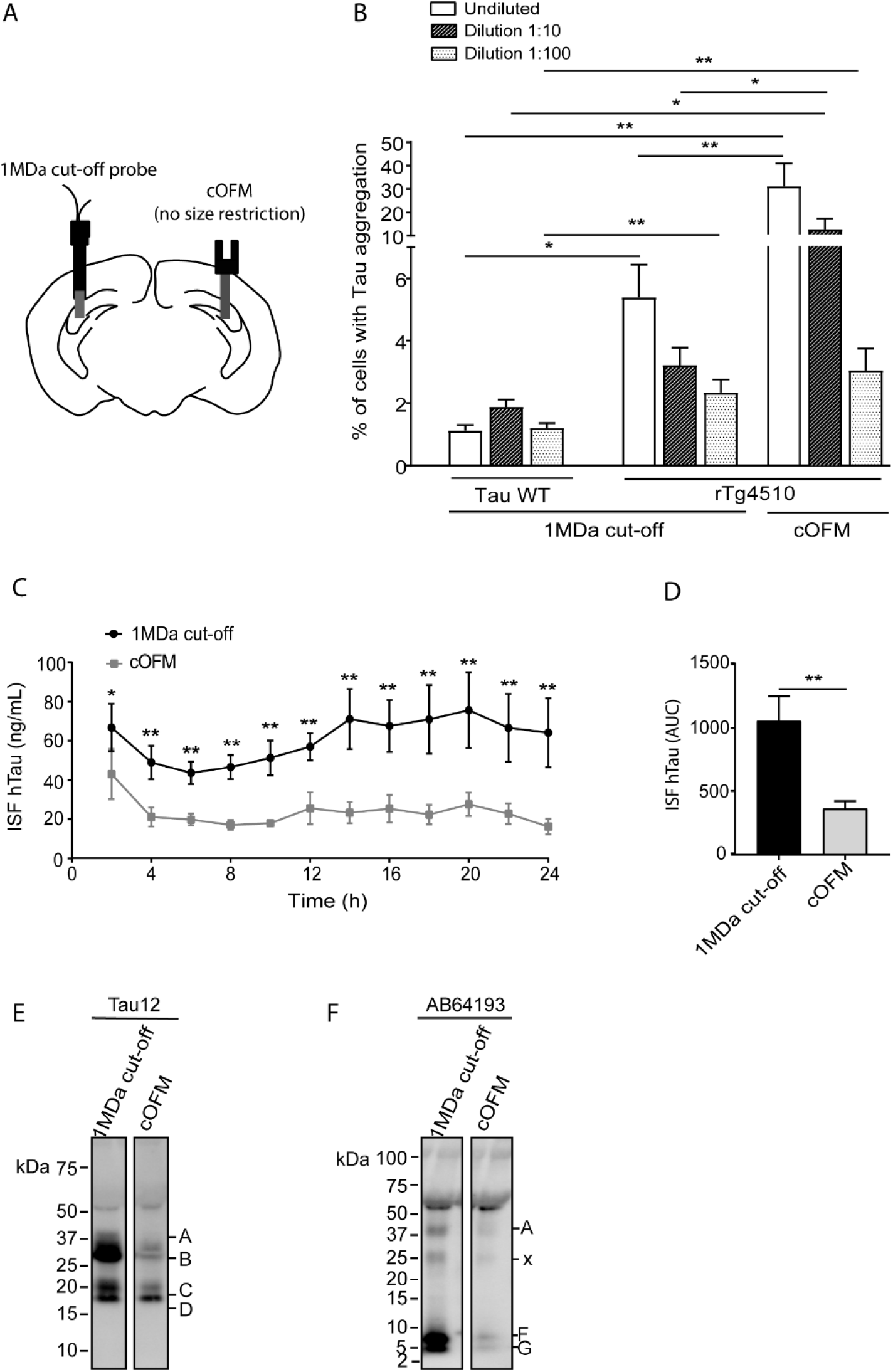
ISF from rTg4510 mice induces Tau aggregation in HEK293-Tau biosensor cells. (A) Schematic of 1MDa cut-off probe and cOFM probe placement. (B) ISF sampled with cOFM or 1MDa cut-off probes from 6-month-old rTg4510 mice was applied to HEK293-Tau biosensor cells undiluted or diluted 10- and 100-fold. (C) hTau levels in ISF sampled by 1MDa cut-off and cOFM probes (p=0.0258). (D) Levels of ISF hTau levels are significantly different between cOFM and 1 MDa cut-off sampling at 4-24 h. Data points represent mean ± SEM. Statistical significance: 2h, p=0.020; 4h, p=0.0012; 6h, p=0.0027; 8h, p=0.0003; 10h, p=0.0002; 12h, p=0.0002; 14h, p<0.0001; 16h, p<0.0001; 18h, p<0.0001; 20h, p<0.0001; 22h, p<0.0001; 24h, p<0.0001. (E) Mean AUC of curves in D (p=0. 0.0008). (F-G) Western blot analysis of Tau fragments in ISF sampled by cOFM and 1 MDa cut-off probes using Tau12 (F) and AB64193 (G) antibodies. Letters A to G indicate Tau fragments (see Figure 2); x indicates non-specific bands. In panels B, C and E, bars represent the mean +SEM. In panel D, lines represent the mean ± SEM. \**p* < 0.05, \*\**p* < 0.01

We next examined whether the distinct seeding propensity of ISF collected by the two probe types was due to differential recovery of Tau or differential sampling of Tau species. To assess hTau recovery, we measured ISF hTau levels from 2 h after probe insertion over 24 h sampling time. hTau levels were significantly higher in ISF sampled with 1MDa cut-off probes than in cOFM ISF (Fig. 3C-D), suggesting that hTau levels are not predictive of seeding propensity. We then investigated the Tau truncation pattern in ISF sampled through cOFM and 1 MDa cut-off probes from 6-month-old rTg4510. The pattern of Tau fragments was comparable in ISF sampled with both probes, as detected by Tau12 (Fig. 3E) and AB64193 antibodies (Fig. 3F), indicating that the sampling procedure does not affect the overall composition of Tau fragments.

Together, these data show that ISF sampled with cOFM displays greater seeding competence compared to ISF obtained with 1MDa cut-off probes, despite containing less hTau and similar Tau fragments. This suggests that the enhanced ability to trigger Tau aggregation may require additional ISF components that are only present after cOFM sampling.

### 3.5. ISF seeding-competence is detectable only in ISF from rTg4510 with Tau inclusions and is independent of Tau fragment content

To investigate ISF seeding competence during the progression of Tau pathology, we sampled ISF from rTg4510 devoid of (2-month-old) and with abundant AT100+ inclusions (6-month-old) using cOFM probes (Fig. 4A). ISF from TauKO mice was also sampled as a control. Only ISF from 6-month-old rTg4510 induced Tau aggregation in HEK293-Tau biosensor cells, while ISF from 2-month-old rTg4510 or TauKO mice did not (Fig. 4A). We therefore investigated whether despite similar truncation patterns (Fig. 2), the levels of specific fragments change with progression of Tau pathology, possibly due to differential aggregation propensity. Levels of Tau fragments in ISF from 2- and 6-month-old rTg4510 were analyzed by western blot (Fig. 4B-E), using Tau12 to detect fragments A, B and C (Fig. 4B), a mix of HT7 and BT2 to identify fragment E (Fig. 4C), AB64193 to analyze fragments F and G (Fig. 4D) and HJ8.3 to detect fragments H, I and J (Fig. 4E). Most ISF Tau fragments (i.e., A, B, C, F, G and J) were significantly more abundant in 2-month-old rTg4510, consistent with the overall greater hTau levels in 2-versus 7-month-old rTg4510 (Fig. 1D).

**Figure 4.**
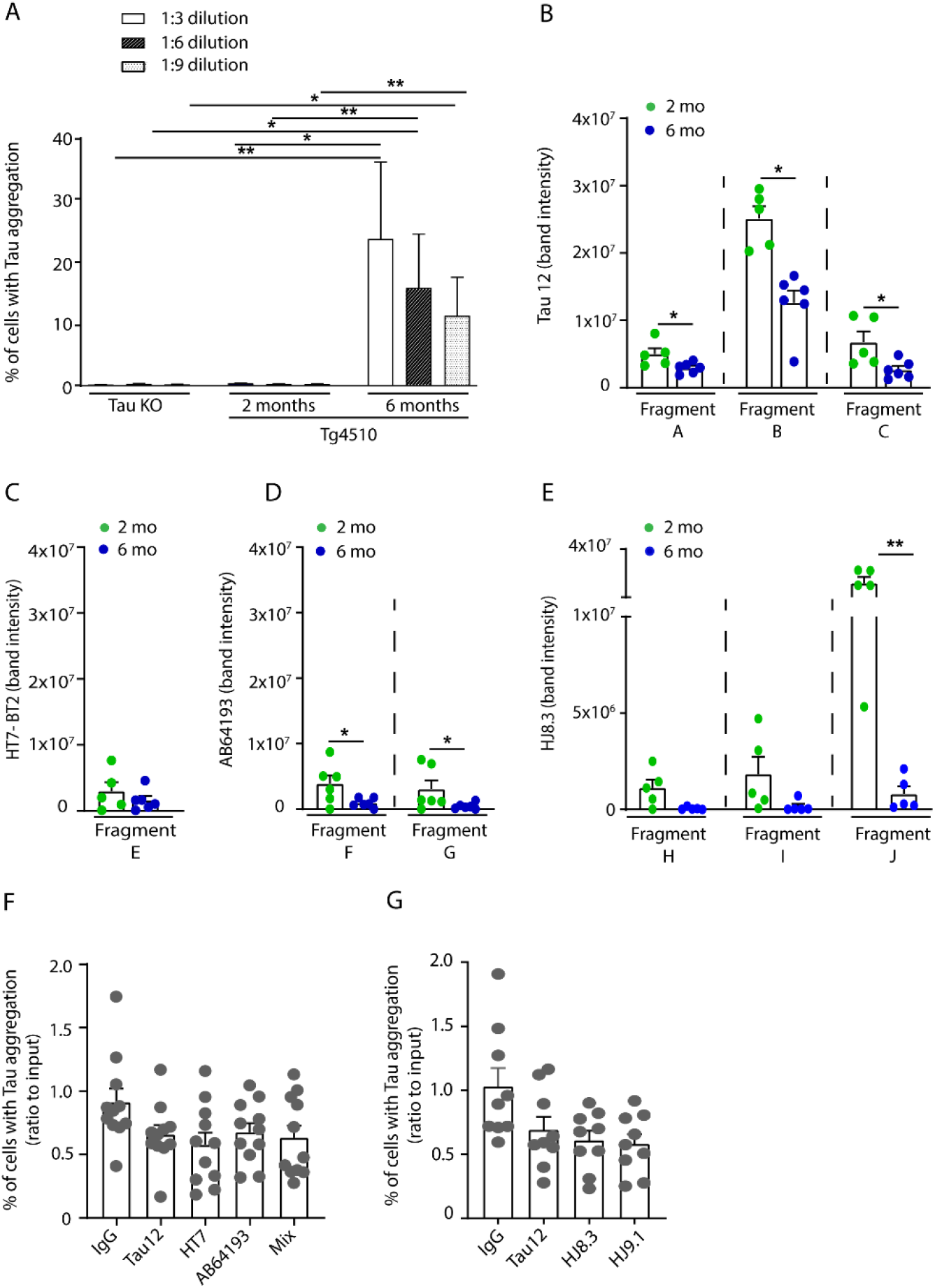
Only ISF from rTg4510 mice with established Tau inclusions induces Tau aggregation in biosensor cells. ISF from rTg4510 and TauKO mice was applied to HEK293-Tau biosensor cells and the formation of Tau aggregates was analyzed as described in the Methods. (A) Seeding competence of ISF from 6-month-old (n=6), 2-month-old rTg4510 (n=9) and TauKO mice (n=3). (B-E) Relative comparison of Tau fragments in ISF from 2-month-old and 7-month-old rTg4510 mice. (B) Fragments A-D detected using Tau12 antibody. Statistical significance: Fragment A, p=0.0472; Fragment B, p=0.0302; Fragment C, p=0.0400. (C) Fragment E detected by the BT2/HT7 antibody mix (p=0.1534). (D) Fragments F-G detected using AB64193 antibody. Statistical significance: Fragment F, p=0.0382; Fragment G, p=0.0420. (E) Fragments H-J detected using HJ8.3 antibody. Statistical significance: Fragment H, p=0.0513; Fragment I, p=0.0613; Fragment J, p=0.0016. (F-G) Immunodepletion of ISF from 6-month-old rTg4510 using Tau12, HT7, AB64193, the mix of the three antibodies (F) or HJ8.3 and HJ9.1 (G). Mouse IgG was used as control in the IP. Statistical comparison vs IgG: (F) Tau12, p=0.2281; HT7, p=0.0767; AB64193, p=0.4841; Mix, p=0.2281; (G) Tau12, p=0.3613; HJ8.3, p=0.1516; HJ9.1, p=0.0781. In A, F and G, bars represent the mean ratio over the input + SEM. In B-E, bars represent the average band intensity + SEM. In B-G, data points indicate values from single mice. In B-C, n=5 for 2- and n=6 for 6-month-old rTg4510; In D, n=6 for 2- and 6-month-old rTg4510; In E, n=5 for 2- and 6-month-old. In F, n=11 and in G, n=9. \**p* < 0.05, \*\**p* < 0.01

Based on these results, we postulated that the overall levels of hTau or specific fragments are not critical for promoting seeding. To test this hypothesis, we immunodepleted Tau fragments using a panel of Tau antibodies spanning the entire Tau sequence (Fig. S4A), including Tau12, HT7 and AB64193 for capturing fragments A-G (Fig. 4F) and HJ9.1 and HJ8.3 for fragments H-J (Fig. 4G). The antibodies were applied either alone or as a mix (Fig. 4F) to ISF from individual 6-month-old rTg4510 mice. Control IgG was used to control for specificity in all experiments (Fig. 4F-G). Immunodepletion with Tau12, AB64193 and HJ8.3 completely removed the respective fragments (Fig. S4B-D) but did not substantially affect ISF seeding competence in HEK293-Tau biosensor cells (Fig. 4F-G). There was a similar lack of change in seeding competence after immunodepletion with HT7, the other mid-domain Tau antibodies BT2 and Tau5 (Fig. S4E) and a C-terminal domain antibody HJ9.1. Control IgG affected neither Tau fragments nor ISF seeding competence.

These results show that seeding competence manifests only in the ISF of mice with abundant inclusions and is independent from the presence of specific Tau fragments.

### 3.6. Phosphorylated Tau is increased in ISF from rTg4510 with Tau inclusions and its removal decreases ISF seeding competence

Analyses of Tau species extracted from brain of rTg4510 mice and AD patients have indicated that seeding-competent Tau species are hyperphosphorylated (Takeda et al., 2016). To investigate the phosphorylation state of ISF Tau, we measured the levels of Tau phosphorylated at sites T181, S198, S199, T231 and S416, which have been reported to be hyperphosphorylated in the soluble brain extracts of early stage AD (i.e., Braak III/IV) (Ercan-Herbst et al., 2019). Levels of Tau phosphorylated at all these sites were significantly higher in ISF from 6-month-old rTg4510 mice than in ISF from 2-month-old mice (Fig. 5A). Using an ELISA based on the phosphorylation- and conformation-dependent antibody AT100 for capture and HT7 as detection, we also found that ISF from 6-month-old rTg4510, but not from TauKO, contains pT212/pS214 Tau (A.U. ± SEM. 6-month-old rTg4510, 21.3 ± 5.6; TauKO, 0.2 ± 0.2).

**Figure 5.**
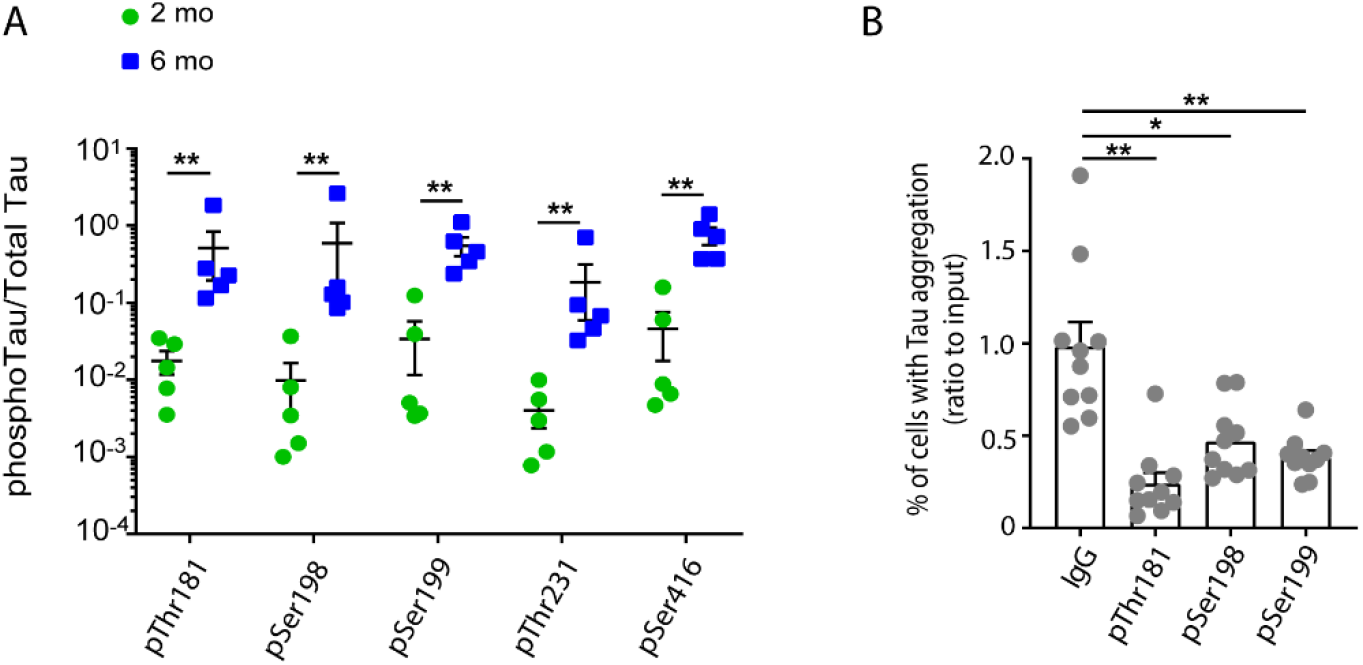
Tau phosphorylation is increased in ISF of rTg4510 mice with Tau inclusions and removal of p-Tau ISF species decreases seeding competence. (A) Phosphorylated Tau in ISF from 2-month-old (n=5) and 6-month-old (n=5) rTg4510 was analyzed as described in the Methods. Phosphorylated Tau values are normalized to total Tau levels measured by a BT2-HT7 MSD assay. Statistical significance: pT181, p=0.0002; pS198, pS199, pT231 and pS416, p<0.0001. (B) Immunodepletion of ISF from 5.3-7-month-old rTg4510 (n=10) using pT181, pS198 or pS199 antibodies. Mouse IgG was used as control in the IP. Statistical comparison vs IgG: pT181, p<0.0001; pS198, p=0.0131; pS199, p=0.0080. Data points indicate values from single mice. In A, lines represent mean values ±SEM. In B, bars represent the mean ratio over the input + SEM. \**p* < 0.05, \*\**p* < 0.01

We next investigated whether removal of phosphorylated Tau reduces ISF ability to seed Tau aggregation in the HEK293 biosensor cells. Immunodepletion of phosphorylated Tau using pT181, pS198 or pS199 Tau antibodies significantly decreased ISF seeding capacity by 76%, 53% or 62%, respectively, as compared to the IgG control (Fig. 5B).

These results indicate that phosphorylated Tau increases in ISF from rTg4510 mice with Tau pathology and is a determinant of ISF seeding propensity.

### 3.7. ISF seeding competence requires aggregated Tau species

To further characterize the ISF species responsible for seeding Tau aggregation, we investigated the potential presence of oligomeric pathological Tau in ISF from 6-month-old rTg4510 mice by ultracentrifugation and analysis of the seeding competence of the resulting pellet and supernatant (Fig. 6A). We found that only the ISF pellet, not the supernatant, promoted Tau aggregation in HEK293-Tau biosensor cells (Fig. 6A). The seeding competence of ISF subfractions did not correlate with hTau concentration. Indeed, levels of hTau in the supernatant were comparable to those in the ISF input and only a small fraction of Tau was detected in the pellet using the MSD hTau assay (Fig. 6B), but not by western blot (Fig S5).

**Figure 6.**
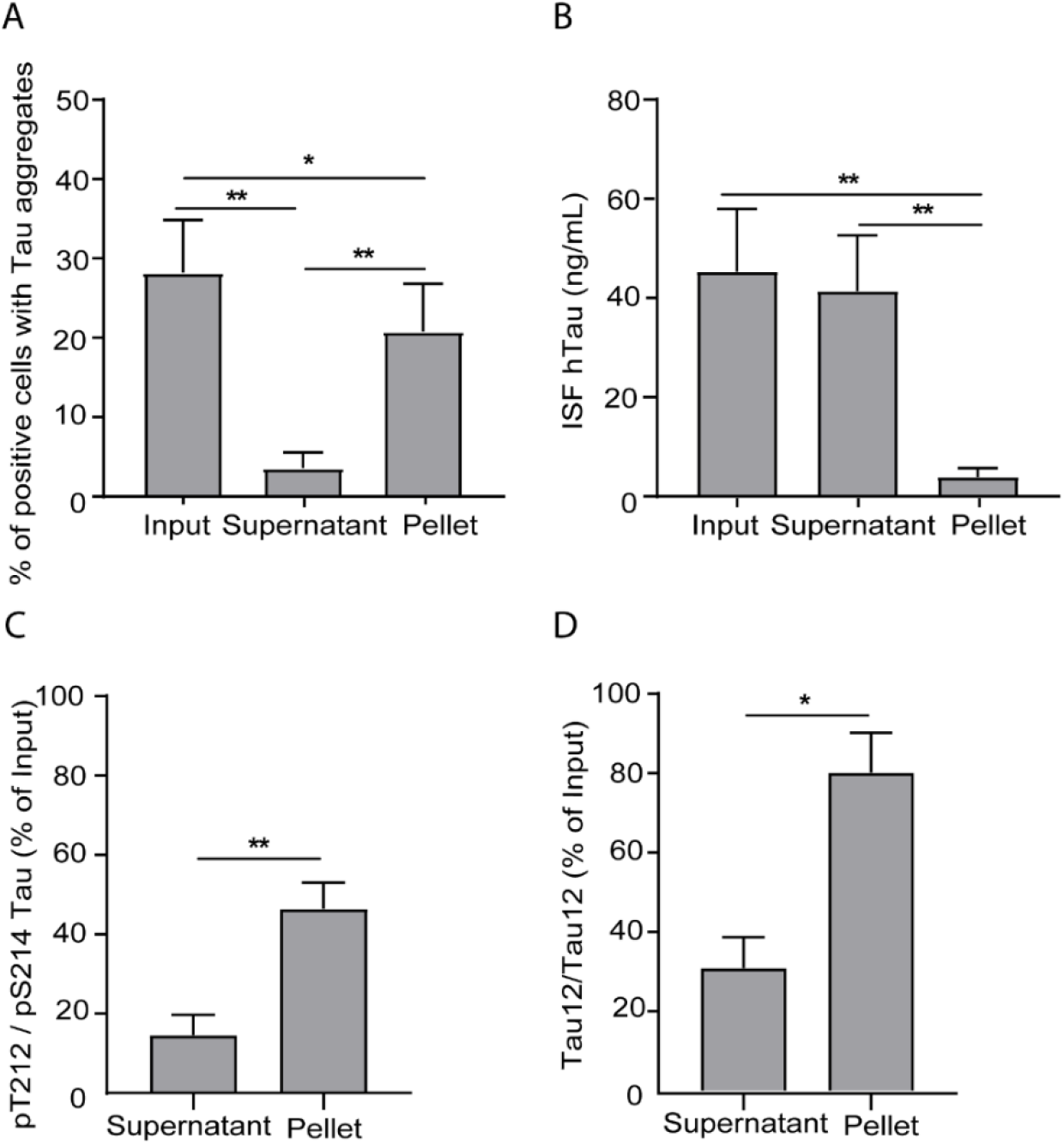
ISF seeding competence requires low-abundance, aggregated and phosphorylated Tau species. ISF of 6-month-old rTg4510 was ultracentrifuged overnight. Seeding competence, levels of hTau and pT212/pS214 Tau in the supernatant and pellet were analyzed as described in the Methods. (A) Seeding competence in HEK293-Tau biosensor cells (n=8). Statistical significance: input vs supernatant, p<0.0001; input vs pellet, p=0.0040; pellet vs supernatant, p=0.0004. (B) hTau levels (n=8). Statistical significance: input vs supernatant, p=0.0868; input vs pellet, p<0.0001; pellet vs supernatant, p=0.0071. (C) pT212/pS214 levels measured by AT100/HT7 Tau ELISA in supernatant and pellet of ISF (p=0.0017). (D) Aggregated Tau detected by Tau/Tau12 symmetric ELISA (n=7) in supernatant and pellet of ISF (n=5, p=0.0187). In graphs C-D, supernatant and pellet were normalized to input. In all graphs, bars represent the mean + SEM. *\*p* < 0.05, \*\**p* < 0.01

Immunoreactivity for AT100, i.e. pT212/pS214 Tau, generally correlates with the presence of pathological Tau aggregates (Augustinack et al., 2002). We thus investigated the distribution of AT100 immunoreactivity in ISF subfractions. pT212/pS214 Tau was significantly elevated in the pellet compared to the supernatant (Fig. 6C). To determine the presence of Tau aggregates, we employed an ELISA assay using the same anti-Tau antibody (Tau12) for capture and detection, that detects only Tau aggregates, e.g. dimer or multimeric forms, but not Tau monomers(Ercan-Herbst et al., 2019). Levels of Tau aggregates were significantly higher in the pellet compared to the supernatant (Fig. 6D).

These results indicate that a small subset of aggregated and/or phosphorylated Tau is present in the ISF of rTg4510 mice bearing Tau inclusions and is likely the driver of ISF seeding competence.

## 4. Discussion

According to the propagation hypothesis, pathological Tau spreads throughout the brain via release and re-uptake between synaptically-connected neurons. Here we report that in freely-moving Tau transgenic mice, hTau is secreted into the ISF in an activity-dependent manner, with dynamics independent of the pathology state, mouse line or brain area. The majority of ISF Tau is truncated, with only a small fraction detected as full-length, and comprises at least 10 distinct fragments spanning the entire Tau protein. The fragmentation pattern is similar across different Tau transgenic models, pathological stages and brain areas. ISF hTau content decreases during Tauopathy progression, whereas its phosphorylation increases. ISF from mice with established Tau inclusions induces Tau aggregation in HEK293-Tau biosensor cells. Removal of phosphorylated Tau, but not Tau fragments, significantly reduces ISF seeding capability indicating a key role of phosphorylated Tau forms in triggering Tau aggregation. Further, ISF seeding-competent Tau is sedimented by ultracentrifugation and constitutes a small fraction of less-soluble Tau enriched in phosphorylated and aggregated forms, which might contribute to the pathological propagation of Tau in vivo.

Previous findings in cultured neurons and in vivo indicate that Tau secretion is an activity-dependent process (Pooler et al., 2013, Sato et al., 2018, Wu et al., 2016, Holth et al., 2019). Indeed, ISF Tau levels vary throughout the wake-sleep cycle according to neuronal activity (Holth et al., 2019) and, in young PS19 transgenic mice, are enhanced by K^+^-evoked neuronal depolarization (Yamada et al., 2011). Here, we confirm and extend previous studies reporting the effect of K^+^-induced depolarization in three different Tau transgenic mouse models at different stages of Tau pathology and in brain areas differently affected by Tau pathology. Consistent with Yamada et al. (2014) findings, we show that K^+^ stimulation transiently increases ISF hTau. This response to K+ depolarization is observed in all three transgenic mouse models, at pre- or post-pathology stages and in different brain areas. Indeed, we find that although basal levels of ISF hTau decrease at advanced stages of Tauopathy, the response to K^+^ stimulation is preserved during the progression of Tau pathology and across different brain areas. In fact, ISF hTau peaks 1 h after K^+^-infusion and returns to baseline levels within 5 h in all Tau transgenic mouse lines. These findings suggest that activity-dependent secretory and homeostatic mechanisms are mostly unaffected by the presence of Tau inclusions or accumulation of phospho-Tau.

In addition, we demonstrate that in Tau transgenic mice, ISF Tau is largely cleaved: only a small fraction of full-length Tau is detectable. At least 10 fragments of variable size spanning the entire Tau sequence are detected, including forms truncated at the N-terminus, C-terminus or both the N- and C-termini. Peptides including only portions of the microtubule binding domain are also present. The fragmentation pattern of ISF Tau is similar in all three transgenic lines investigated and is constant during Tauopathy progression or and across brain areas. These novel data extend and reconcile previous inconsistent findings reporting either full-length Tau or a 25 kDa truncated Tau form as prominent Tau species in ISF of PS19 and JNPL3 mice, respectively (Yamada et al., 2014, Bright et al., 2015) and indicate that ISF Tau comprise a complex mix of proteoforms. The origin of ISF cleaved forms remains unclear. Tau cleavage may occur intraneuronally before release, in the extracellular space upon secretion or after uptake and processing by microglia (Behrendt et al., 2019, Bolos et al., 2016, Asai et al., 2015, Maphis et al., 2015). The observation that cultured neurons in vitro release truncated forms of Tau (Kanmert et al., 2015) stretching the entire Tau sequence (Sato et al. 2018), suggests that neurons are both required and sufficient to generate extracellular Tau fragments. Further studies are warranted to investigate non-cell autonomous mechanisms of Tau truncation in the brain of living mice.

Although we find a constant truncation pattern in all three Tau transgenic lines, the overall amount of ISF hTau is reduced as pathology progresses, consistent with previous results in PS19 mice (Yamada et al., 2011). This effect appears to be associated with pathological changes in Tau species rather than aging. In fact, ISF levels of endogenous murine Tau do not significantly change with age in either wild type C57Bl6 or rTg4510 transgenic mice. In contrast to the overall Tau amount, ISF levels of phospho-Tau are elevated in rTg4510 mice with established Tau inclusions: phosphorylation is increased at Tau residues hyperphosphorylated in AD brain at early Braak stages, such as pT181, pS198, pS199, pT231 and pS416 (Ercan-Herbst et al., 2019). Moreover, ISF Tau phosphorylated at T212/S214 is elevated as detected by the phosphorylation- and conformation-dependent antibody AT100. Although it is well known that Tau is hyperphosphorylated in AD and transgenic mouse brain, this is the first study to reveal phosphorylated Tau species in ISF.

Phospho-Tau appears a key driver of ISF seeding competence. Indeed, removal of phospho-Tau using antibodies recognizing phospho-sites detected in early AD (i.e., pT181, pS198 or pS199) significantly decreases the ability of ISF to trigger Tau aggregation in the HEK293-Tau biosensor cells. Previous findings indicate that pathological, seeding-competent Tau species from AD and Tg4510 mouse brain are hyperphosphorylated (Takeda et al., 2015): we now demonstrate that phospho-Tau is secreted into ISF, retaining seeding competence. In contrast, depletion of soluble ISF Tau fragments with antibodies against unmodified Tau does not significantly affect ISF seeding propensity. This finding matches to the observation that ISF Tau fragment levels are elevated in rTg4510 mice prior to pathology onset, when ISF from these mice is not able to induce Tau aggregation.

Our findings also indicate that ISF seeding competent Tau comprises a subset of less soluble Tau species that can be separated by ultracentrifugation and are enriched in phosphorylated and aggregated forms. Previous findings showed that Tg4510 ISF Tau is taken up by neurons in cultures (Takeda et al., 2015) and can induce Tau aggregation in HEK293 biosensor cells (Takeda et al., 2016). We now report for the first time that the less soluble, aggregated ISF Tau found in the ultracentrifugation pellet are responsible for promoting Tau aggregation in biosensor cells. This observation is consistent with previously reported high-molecular-weight seeding-competent Tau species isolated by size exclusion chromatography from brain of 12-month-old rTg4510 mice and AD brain tissue (Takeda et al., 2015). The minute amount of ISF seeding-competent species poses significant technical challenges to their investigation. Additional efforts are warranted to further clarify the nature of Tau seeding-competent species, their aggregation state, the presence of exosomes and/or interacting partners.

There is consistent evidence that the transneuronal propagation of pathological Tau underlies the progression of Tauopathy in AD. Blocking Tau spreading using Tau antibodies therefore represents an attractive therapeutic strategy for AD, which is currently being pursued by several groups worldwide (Bright et al., 2015, Yanamandra et al., 2015, Albert et al., 2019, Rosenqvist et al., 2018, Vandermeeren et al., 2018). Here we demonstrate for the first time that in Tau transgenic mice, ISF Tau is comprises multiple distinct truncated and phosphorylated forms. Furthermore, we show that a small subset of less soluble, aggregated and phosphorylated Tau species is capable of inducing Tau assembly in biosensor cells in vitro. Considering their potential implication in the progression of Tau pathology in AD and their impact in informing the discovery of new AD therapeutics, further investigation is warranted to identify Tau propagating species.

## Author contributions

JH, SY, FP, IM, MWM and EB performed microdialysis experiments; EB, GP and ERC performed biochemical analysis; EB, GP, ERC and SB performed in vitro seeding experiments; YM performed statistical analysis; AS, EEH and DEE performed phospho-Tau and aggregated Tau analyses; CK and KB performed histological analysis; MM and MC supported data analysis and discussion and interpretation of the results; EB, LG and KS designed and coordinated experiments and data analyses; EB and LG wrote the manuscript and EB, LG and KS reviewed and edited the manuscript. All authors critically discussed the results, reviewed and approved the manuscript.

## Disclosure

EB, GP, SJ, YM, FP, IM, MM, SB, MC, AS, DEE, KB, CK, LG, KS are employees of AbbVie. JH, ERC, MWM were employees of AbbVie at the time of the study. DEE, EH were employees of BioMedX, Heidelberg, working in a team sponsored by AbbVie during the time of the study. The design, study conduct and financial support for this research were provided by AbbVie. AbbVie participated in the interpretation of data, review, and approval of the publication.

This project has received funding from the Innovative Medicines Initiative 2 Joint Undertaking under grant agreement No. 116060 (IMPRiND). This Joint Undertaking receives support from the European Union’s Horizon 2020 research and innovation programme and EFPIA. This work is supported by the Swiss State Secretariat for Education, Research and Innovation (SERI) under contract number 17.00038. The opinions expressed and arguments employed herein do not necessarily reflect the official views of these funding bodies.

## Acknowledgements

We would like to thank Thomas Altendorfer-Kroath and Thomas Birngruber from JOANNEUM RESEARCH for providing cerebral Open-Flow Microperfusion (cOFM) probes. No funding to disclose. Special thanks to Maria Vasileva-Diehl of Abbvie for helping in the publication process and to Diana Clausznitzer of AbbVie for the interesting and valuable scientific discussions.

## Abbreviations

aCSF: artificial cerebrospinal fluid
AD: Alzheimer disease
CBD: corticobasal degeneration
cOFM: cerebral open flow microperfusion
FTD: frontotemporal dementia
hTau: human Tau
ISF: interstitial fluid
IP: immunoprecipitation
mTau: mouse Tau
PSP: progressive parasupranuclear palsy
TauKOTau: knockout mouse line

