## Supplementary figures for "Tau in the brain interstitial fluid is fragmented and seeding-competent"

### Supplementary tables and figures

### Supplementary Table 1: Statistical analyses

| *Figure* | *Data displayed as* | *Normality* | *Transformation* | *Homogeneity of variances* | *Statistical method used* |
|---|---|---|---|---|---|
| *1A* | *Scatterplot with bar: mean with SEM* | *Normality per Shapiro-Wilk test after transformation. 4m.o.: p=0.8240, 9m.o.: p=0.5260* | *Log transformation* | *Yes (Levene's p=0.2828)* | *Two-sample t-test* |
| *1B* | *Mean ± SEM* | *Normality per Shapiro-Wilk test, p-values for each timepoint: 1h: K+100mM: 0.2431, veh: 0.7968; 2h: K+100mM: 0.7881, veh: 0.8917; 3h: K+100mM: 0.0592, veh: 0.1777; 4h: K+100mM: 0.5616, veh: 0.6041; 5h: K+100mM: 0.3720, veh: 0.7897; 6h: K+100mM: 0.8825, veh: 0.2753; 7h: K+100mM: 0.2564, veh: 0.2259* | *No* | *Yes (Levene's p=0.0960)* | *Mixed effects model followed by test contrast* |
| *1C* | *Scatterplot: mean with SEM* | --- | *No* | --- | *Non-param. Fisher-Pitman for paired observations* |
| *1D* | *Scatterplot with bar: mean with SEM* | *Normality per Shapiro-Wilk test after transformation. 2m.o.: p=0.1303, 7m.o.: p=0.2268* | *Log transformation* | *Yes (Levene's p=0.6986)* | *Two-sample t-test* |
| *1E* | *Mean ± SEM* | *Normality of the distribution of differences between paires per Shapiro-Wilk test for the following timepoints: 3h:0.7839; 4h: 0.5361, 5h: 0.7279; rejected normality per Shapiro-Wilk test: 1h , 2h* | *Log transformation* | *No overall homogeneity of variances, but homogeneity of variances at each timepoint (Levene's p-value) at 1h: 0.8845, 2h: 0.6906, 3h: 0.1809, 4h: 0.1361, 5h: 0.0915)* | *Per timepoint: Paired t-test and Wilcoxon test* |
| *1F* | *Scatterplot: mean with SEM* | *Normality of the distribution of differences between paires per Shapiro-Wilk test: p=0.5555* | *No* | *Yes (Levene´s p=0.1341* | *Paired t-test* |
| *1G* | *Scatterplot with bar: mean with SEM* | *Normality of the distribution of differences between pairs after transformation, Shapiro-Wilk p=0.4194* | *Log transformation* | *Yes (Levene's p=0.1130)* | *Paired t-test* |

<table>
<tr><td>1H</td><td>Mean ± SEM</td><td>Normality per Shapiro-Wilk test, p-values for each timepoint: 1h: cortex: 0.6042, hippo: 0.5557; 2h: cortex: 0.7536, hippo: 0.8273; 3h: cortex: 0.4138, hippo: 0.6899; 4h: cortex: 0.6768, hippo: 0.2886; 5h: cortex: 0.2632, hippo: 0.3048; 6h: cortex: 0.3251, hippo: 0.8537; 7h: cortex: 0.5443, hippo: 0.5871</td><td>No</td><td>Yes (Levene's p=0.08335)</td><td>Mixed effects model followed by test contrast</td></tr>
<tr><td>3B</td><td>Bars: mean with SEM</td><td>Normality per Shapiro-Wilk test for each group each dilution, p-values. For pure: Tau WT ISF 0.9999, Tg4510 ISF 1MDa 0.9291, Tg4510 ISF Open Flow 0.4635. For 1:10: Tau WT ISF 0.9999, Tg4510 ISF 1MDa 0.3225, Tg4510 ISF Open Flow 0.0584. For 1:100: Tau WT ISF 0.9999, Tg4510 ISF 1MDa 0.9718, Tg4510 ISF Open Flow 0.4856</td><td>Log transformation</td><td>Yes (Levene's p for pure: 0.0933, for 1:10: 0.4255; for 1:100: 0.0697)</td><td>One-way ANOVA (least squares fit for each dilution) followed by Tukey's HSD test<br><br>Significant results:<br><table>
<tr><th>Level</th><th>-Level</th><th>p-values</th></tr>
<tr><td>Tg4510 ISF cOFM 1:10</td><td>Tau WT ISF 1:10</td><td>0.0131*</td></tr>
<tr><td>Tg4510 ISF cOFM 1:10</td><td>Tg4510 ISF 1MDa 1:10</td><td>0.0195*</td></tr>
<tr><td>Tg4510 ISF cOFM 1:100</td><td>Tau WT ISF 1:100</td><td>0.0021*</td></tr>
<tr><td>Tg4510 ISF 1MDa 1:100</td><td>Tau WT ISF 1:100</td><td>0.0074*</td></tr>
<tr><td>Tg4510 ISF cOFM pure</td><td>Tau WT ISF pure</td><td>0.0003*</td></tr>
<tr><td>Tg4510 ISF cOFM pure</td><td>Tg4510 ISF 1MDa pure</td><td>0.0026*</td></tr>
<tr><td>Tg4510 ISF 1MDa pure</td><td>Tau WT ISF pure</td><td>0.0360*</td></tr>
</table></td></tr>
<tr><td>3C</td><td>Mean ± SEM</td><td>Normality per Shapiro-Wilk test after transformation, p-values for each time point, each group. 2h: conv, 0.0629, cOFM, 0.9185; 4h: conv, 0.0550, cOFM, 0.9847; 6h: conv, 0.0651, cOFM, 0.1544; 8h: conv, 0.1084, cOFM, 0.9581; 10h: conv, 0.1460, cOFM, 0.8370; 12h: conv, 0.4352, cOFM, 0.2087; 14h: conv, 0.5078, cOFM, 0.3067; 16h: conv, 0.7198, cOFM, 0.4510; 18h: conv, 0.8739, cOFM, 0.6409; 20h: conv, 0.1719, cOFM, 0.0892; 22h: conv, 0.5585, cOFM, 0.2094; 24h: conv, 0.5777, cOFM, 0.1033</td><td>Cube root</td><td>Yes (Levene's p=0.1090)</td><td>Mixed effects model followed by test contrast</td></tr>
<tr><td>3D</td><td>Bars: mean with SEM</td><td>Normality of the distribution of differences between paires after transformation, Shapiro-Wilk p=0.0501</td><td>Log transformation</td><td>Yes (Levene's p=0.6660)</td><td>Paired t-test</td></tr>
<tr><td>4A</td><td>Bars: mean with SEM</td><td>---</td><td>---</td><td>---</td><td>Non-parametric each-pair Wilcoxon and all-pairs Dunn's test<br><br>Significant results:<br><table>
<tr><th>Level</th><th>Level</th><th>p-value</th></tr>
<tr><td>6 months 1:3</td><td>KO 1:3</td><td>0.0016*</td></tr>
<tr><td>2 months 1:3</td><td>6 months 1:3</td><td>0.0251*</td></tr>
<tr><td>6 months 1:6</td><td>KO 1:6</td><td>0.0282*</td></tr>
<tr><td>2 months 1:6</td><td>6 months 1:6</td><td>0.0018*</td></tr>
</table></td></tr>
</table>

| | | | | | |
|---|---|---|---|---|---|
| | | | | | 6 months 1:9 / KO 1:9 / 0.0282*<br>2 months 1:9 / Old 1:9 / 0.0018* |
| 4B | Scatterplot with bar: mean with SEM | Normality per Shapiro-Wilk test after transformation for each fragment, each group: p-values. Fragm. A: 6m.o. 0.7711, 2m.o. 0.5875. Fragm. B: 6m.o. 0.4200, 2m.o. 0.2208. Fragm. C: 6m.o. 0.9785, 2m.o. 0.2614. | Log transformation | Yes (Levene's p-value for fragm. A: 0.3560; fragm. B: 03239; fragm. C: 0.7994. | Two-sample t-test for each fragment |
| 4C | Scatterplot with bar: mean with SEM | Normality per Shapiro-Wilk test after transformation, p-values for 2m.o.: 0.3491, for 6m.o.: 0.0540 | Log transformation | Yes (Levene's p=0.7305) | Two-sample t-test |
| 4D | Scatterplot with bar: mean with SEM | Normality per Shapiro-Wilk test for each fragment, each group: p-values. Fragm. F: 6m.o. 0.2786, 2m.o. 0.3136. Fragm. G: 6m.o. 0.1975<br><br>Normality rejected fragm. G: 2m.o. 0.0036 | No | No (Levene's p-value for fragm. F: 0.0279; for fragm. G: 0.0149) | Welch's test for fragm. F and Kruskal-Wallis test for fragm. G |
| 4E | Scatterplot with bar: mean with SEM | Normality per Shapiro-Wilk test after transformation for each fragment, p-values. Fragm. H: 6m.o. 0.8726, 2m.o. 0.3387. Fragm. I: 6m.o. 0.7726, 2m.o. 0.9098. Fragm. J: 6m.o. 0.5021, 2m.o. 0.1081 | Log transformation | Yes (Levene's p-value for fragm. H: 0.3308; fragm. I: 0.8142; fragm. J: 0.1356) | Two-sample t-test for each fragment |
| 4F | Scatterplot with bar: mean with SEM | --- | --- | ---- | Non-parametric Dunn's test |
| 4G | Scatterplot with bar: mean with SEM | --- | --- | ---- | Non-parametric Dunn's test |
| 5A | Scatterplot: mean ± SEM | Normality per Shapiro-Wilk test after transformation for each group, p-values. pT181: 6m.o. 0.1510, 2m.o. 0.6708. pS198: 6m.o. 0.1240, 2m.o. 0.7131. pS199: 6m.o. 0.9801, 2m.o. 0.1352. pT231: 6m.o. 0.1796, 2m.o. 0.7852. pS416: 6m.o. 0.4258, 2m.o. 0.2609 | Log transformation | Yes (Levene's p-value 0.2191) | Two way ANOVA with interaction followed by test contrast |
| 5B | Scatterplot with bar: mean with SEM | --- | --- | ---- | Non-parametric Dunn's test |
| 6A | Bars: mean with SEM | Normality of the distribution of differences between paires per | Sq. root/No transformation | Yes (Levene's p-values: 0.0922, | Paired t-tests |

| | | | | | |
|---|---|---|---|---|---|
| | | *Shapiro-Wilk test , p-values: 0.0540, 0.1294, 0.5199* | | *0.6167, 0.4727)* | |
| *6B* | *Bars: mean with SEM* | *Normality of the distribution of differences between pairs per Shapiro-Wilk test after transformation, p-values: 0.0896, 0.0565, 0.5014* | *Log transformation* | *Yes (Levene's p-values: 0.2510, 0.2636, 0.9843)* | *Paired t-tests* |
| *6C* | *Bars: mean with SEM* | *Normality of the distribution of differences between pairs per Shapiro-Wilk test after transformation, p-value: 0.5654* | *Log transformation* | *Yes (Levene's p-value: 0.4109)* | *Paired t-test* |
| *6D* | *Bars: mean with SEM* | *Normality of the distribution of differences between pairs per Shapiro-Wilk test after transformation, p-value: 0.5100* | *Log transformation* | *Yes (Levene's p-value: 0.5401)* | *Paired t-test* |
| *S2A* | *Mean ± SEM* | *Normality per Shapiro-Wilk test, p-values for each time point, each group. 1h: K+100mM, 0.8466, veh, 0.8920; 2h: K+100mM, 0.2645, veh, 0.8290; 3h: K+100mM, 0.8749, veh, 0.9848; 4h: K+100mM, 0.7637, veh, 0.3115; 5h: K+100mM, 0.9786, veh, 0.2458* | *No* | *Yes (Levene's p=0.1704)* | *Mixed effects model followed by test contrast* |
| *S2B* | *Scatterplot with bar: mean with SEM* | *Normality of the distribution of differences between pairs per Shapiro-Wilk test, p-value: 0.2708* | *No* | *Yes (Levene's p-value 0.1339)* | *Paired t-test* |
| *S2 C* | *Mean ± SEM* | *Normality per Shapiro-Wilk test after transformation, p-values for each time point, each group. 1h: cortex, 0.0561, hippo, 0.2046; 2h: cortex, 0.5127, hippo, 0.6287; 3h: cortex, 0.0863, hippo, 0.9957; 4h: cortex, 0.3011, hippo, 0.1423; 5h: cortex, 0.3116, hippo, 0.5833; 6h: cortex, 0.2094, hippo, 0.9830; 7h: cortex, 0.0730, hippo: 0.6523* | *Log transformation* | *Yes (Levene's p-value 0.1525)* | *Mixed effects model followed by test contrast* |
| *S4 E* | *Scatterplot with bar: mean with SEM* | *Normality per Shapiro-Wilk test after transformation, p-values for each group. IgG IP 0.2891; Tau12 IP 0.7723; HT7 0.2504; BT2 0.8189* | *Log transformation* | *Yes (Levene's p-value 0.0844)* | *Dunnett's test* |

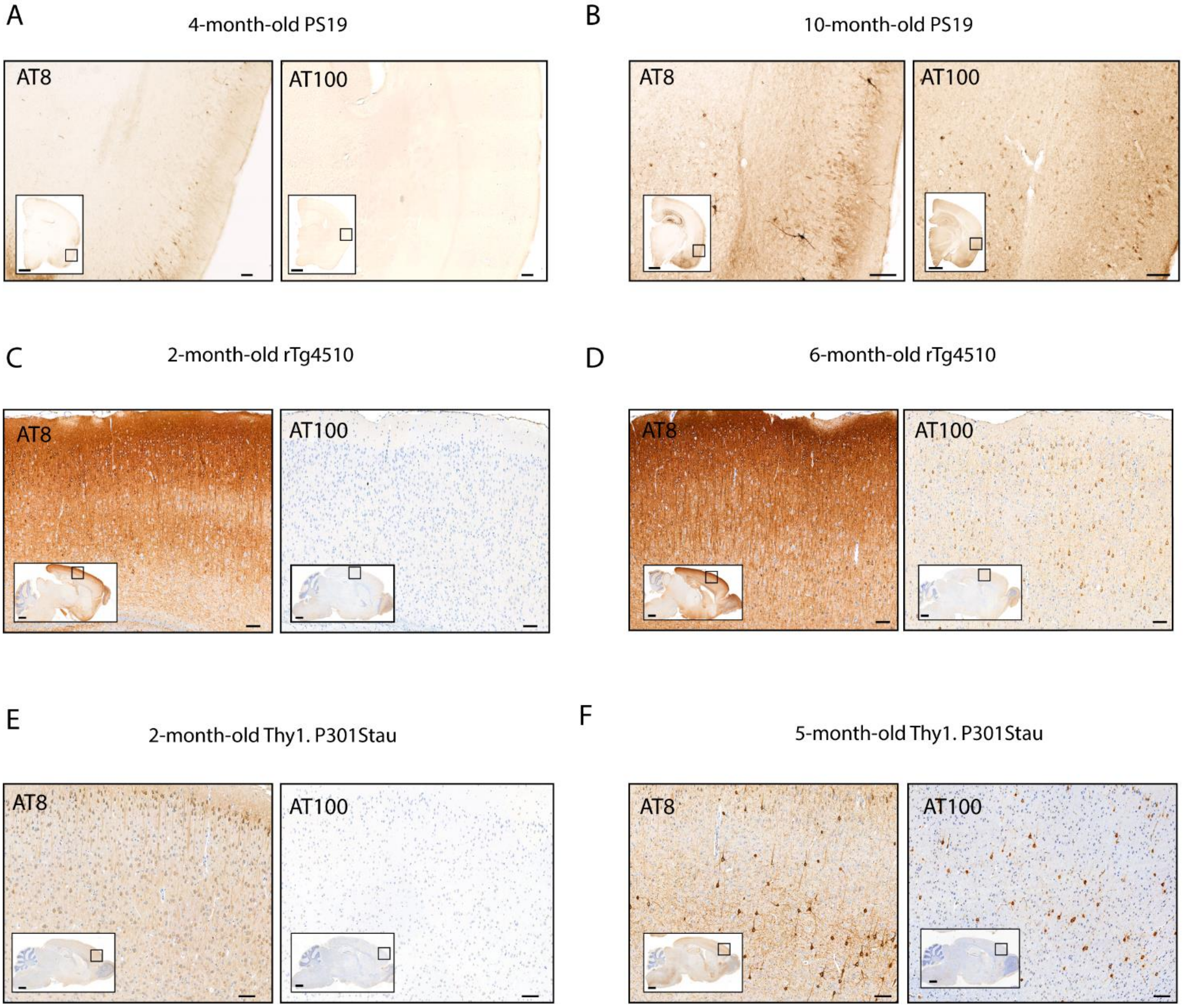


**Figure S1. Tau pathology in Tau transgenic mice**

Immunoreactivity for AT8 and AT100 Tau antibodies in the cortex of PS19 (A-B), rTg4510 (C-D) and Thy1.P301Stau (E-F) at various ages. In C-F, nuclei are counterstained by NeuN. Whole brain coronal or sagittal sections are shown in insets. Square boxes in insets indicate the area shown at higher magnification in the respective main panel. Scale bars: 100 μm in main panels and 1000 μm in insets.

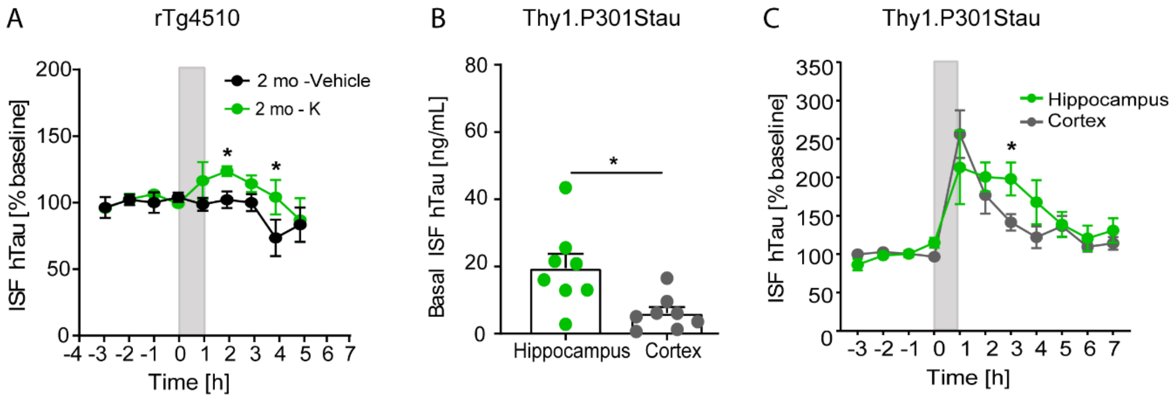


**Figure S2. ISF hTau levels in basal conditions or upon K$^+$ stimulation in 2-month-old rTg4510 and 2-month-old Thy1.P301Stau mice**

(A) ISF hTau in K$^+$- or vehicle-perfused 2-month-old rTg4510 mice (n=4). Statistical significance: 1 h, p=0.1422; 2h, p=0.0490; 3h, p=0.2383; 4h, p=0.0308; 5h, p=0.7681. (B) Basal levels of ISF hTau in hippocampus and cortex of Thy1.P301Stau mice (n=8; p=0.0131). (C) ISF hTau levels in the cortex and hippocampus of Thy1.P301Stau. Grey shading indicates the period of perfusion with K$^+$ or vehicle. ISF hTau levels are expressed as a percentage of baseline. Statistical significance: 1h, p=0.1105; 2h, p=0.2801; 3h, p=0.0471; 4h, p=0.0752; 5h, p=0.9256; 6h, p=0.7706; 7h, p=0.5308. In A and C, ISF hTau is expressed as mean % of baseline ± SEM. In B, bars represent the mean + SEM and data points indicate values from single mice. *$p < 0.05$

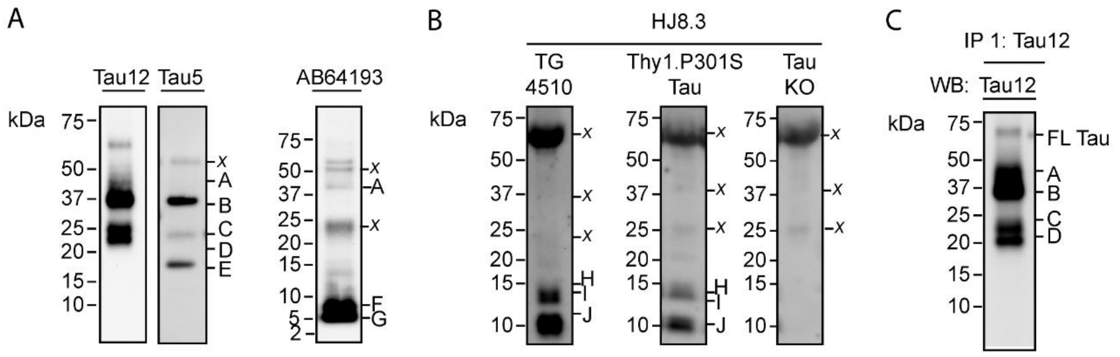


**Figure S3. Tau is truncated in the ISF of Tau transgenic mice**

Pooled ISF fractions from Tau transgenic mice were immunoprecipitated using Tau12, HJ8.3 or the antibody mix Tau12, HT7, AB64193 and HJ9.1 and analyzed by western blot as described in the Methods. (A) Tau fragments in ISF from 2-month and 9-month-old PS19 mice. (B) Tau fragments in ISF from 6-month-old rTg4510 mice and 5-month-old Thy1.P301Stau mice. TauKO ISF was used as a negative control. (C) Full-length Tau (FL Tau) was detected by western blot with Tau12 at longer exposure times. Capital letters (A-J) indicate Tau fragments identified in ISF by Tau12. x indicates non-specific bands.

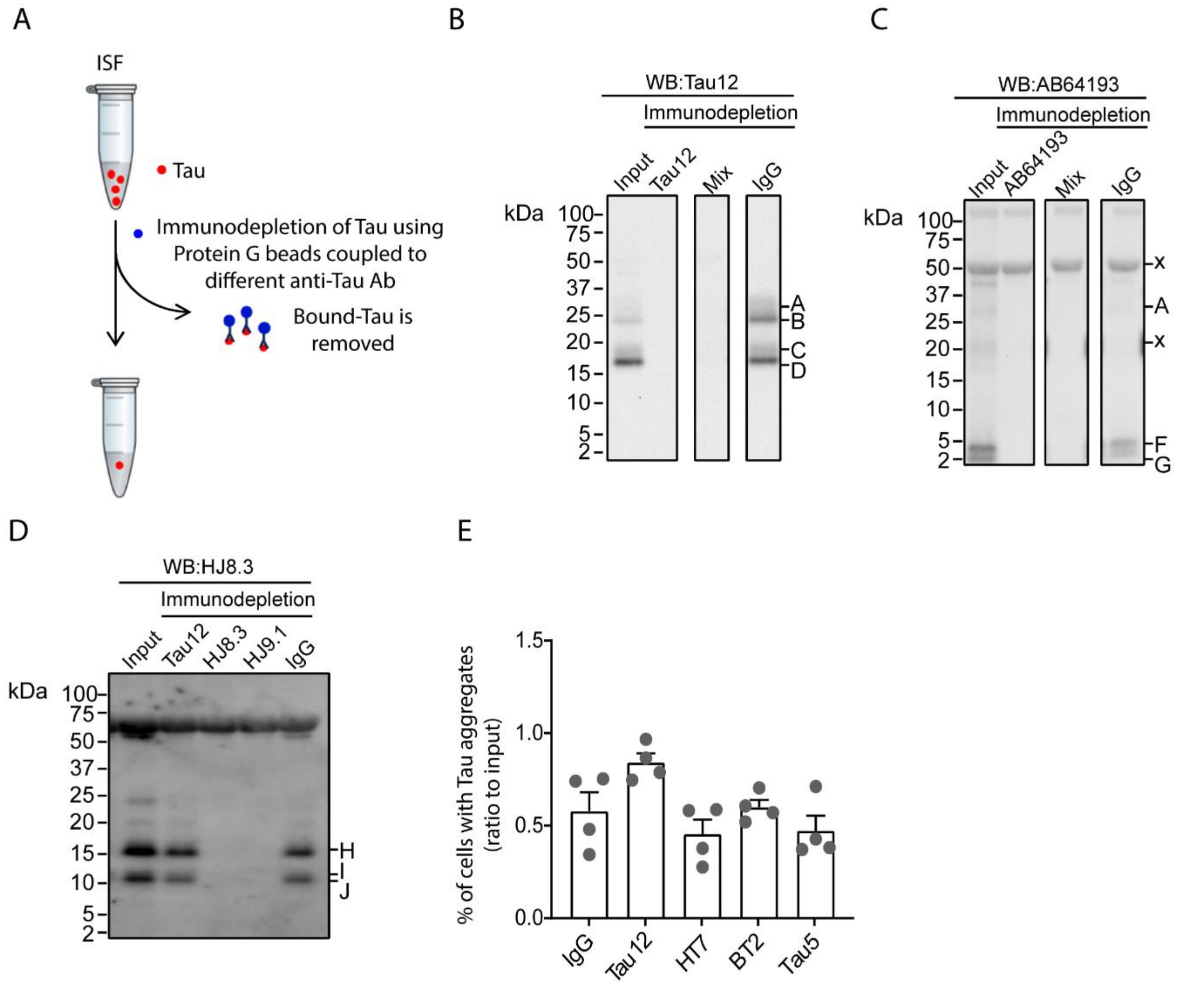


**Figure S4. Immunodepletion of Tau fragments with a panel of antibodies for unmodified Tau does not significantly affect ISF seeding competence**

ISF samples were immunodepleted using Tau antibodies and analyzed by western blot as described in the Methods. (A) Schematic of the experimental procedure. (B-D) Western blot analysis of Tau in immunodepleted samples and input. Antibodies used for IP and western blot are indicated in the respective panels. In panels B-C, the antibody mix Tau12, HT7, AB64193 and HJ9.1 (Mix) was also used for IP. In panels B-D, capital letters A-J indicate Tau fragments identified in ISF. x indicates non-specific bands. Mouse IgG was used as control for the IP. (E) Seeding competence in ISF from 6-month-old rTg4510 (n=4) after immunodepletion using Tau antibodies Tau12, HT7, BT2 and Tau5 or the mouse control IgG. Data are expressed as a ratio over the seeding competence of the input. Statistical comparison vs IgG: Tau12, p=0.0695; BT2, p=0.9987; Tau5, p=0.6954; HT7, p=0.5842. In panel E, bars represent mean + SEM.

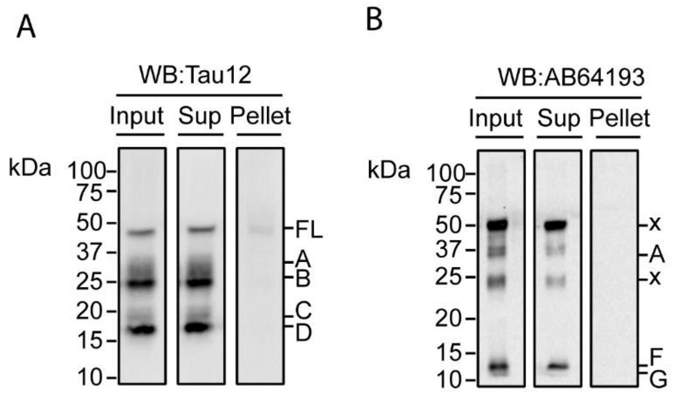


**Figure S5. Tau fragments are detectable by western blot in ISF supernatant but not in the pellet**

Pooled ISF fractions were fractionated by ultracentrifugation and Tau fragments in supernatant, pellet and input were analyzed by western blot using Tau12 (A) and AB64193 (B) antibodies. Capital letters A-G indicate Tau fragments identified in ISF. x indicates non-specific bands.
